# Robust prediction of relative binding energies for protein-protein complex mutations using free energy perturbation calculations

**DOI:** 10.1101/2024.04.22.590325

**Authors:** Jared M. Sampson, Daniel A. Cannon, Jianxin Duan, Jordan C. K. Epstein, Alina P. Sergeeva, Phinikoula S. Katsamba, Seetha M. Mannepalli, Fabiana A. Bahna, Hélène Adihou, Stéphanie M. Guéret, Ranganath Gopalakrishnan, Stefan Geschwindner, D. Gareth Rees, Anna Sigurdardottir, Trevor Wilkinson, Roger B. Dodd, Leonardo De Maria, Juan Carlos Mobarec, Lawrence Shapiro, Barry Honig, Andrew Buchanan, Richard A. Friesner, Lingle Wang

**Author notes:** Corresponding author *Email addresses:* (Richard A. Friesner), (Lingle Wang). Current address: Lundbeck, Copenhagen, Denmark (Jianxin Duan). Current address: Research and Development, Euroapi France, Paris, France (Hélène Adihou).

## Abstract

Computational free energy-based methods have the potential to significantly improve throughput and decrease costs of protein design efforts. Such methods must reach a high level of reliability, accuracy, and automation to be effectively deployed in practical industrial settings in a way that impacts protein design projects. Here, we present a benchmark study for the calculation of relative changes in protein-protein binding affinity for single point mutations across a variety of systems from the literature, using free energy perturbation (FEP+) calculations. We describe a method for robust treatment of alternate protonation states for titratable amino acids, which yields improved correlation with and reduced error compared to experimental binding free energies. Following careful analysis of the largest outlier cases in our dataset, we assess limitations of the default FEP+ protocols and introduce an automated script which identifies probable outlier cases that may require additional scrutiny and calculates an empirical correction for a subset of charge-related outliers. Through a series of three additional case study systems, we discuss how protein FEP+ can be applied to real-world protein design projects, and suggest areas of further study.

**Graphical Abstract:** 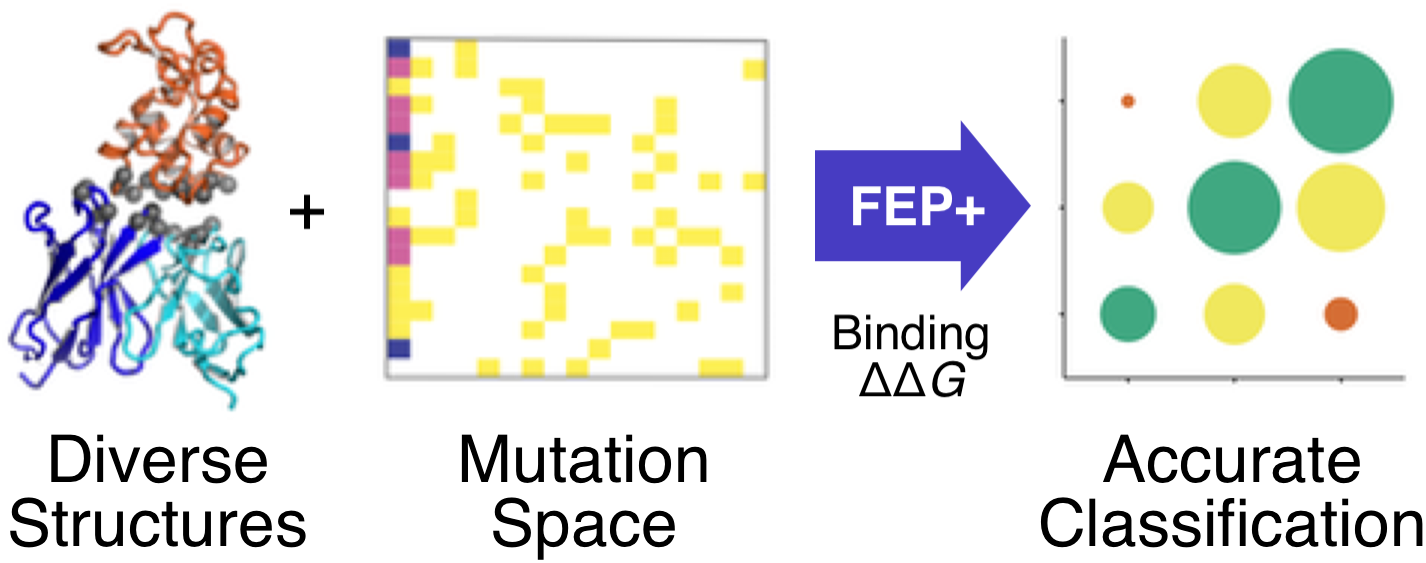

**Highlights:**

- Reliable calculation of relative binding free energy changes for most protein mutations to within ∼1 kcal/mol.
- Automated Protein FEP+ Groups treatment of alternate protonation states for titratable residues.
- Application of FEP+ methodology to “real-world” protein design projects.

## Introduction

Optimization of protein-protein binding interfaces is central to biologics design. Such interfaces can be altered by protein residue mutation to increase or decrease binding affinity, as well as other properties such as thermostability, hydrophobicity, immunogenicity, and pH sensitivity [1, 2, 3, 4, 5]. Traditionally, this has been done through a combination of chemical and structural intuition on the part of research scientists and large-scale experimentation, via such methods as brute-force saturation mutagenesis followed by a high-throughput assay; random or degenerate codon mutagenesis; or library screening using a display technology [6, 7, 8, 9]. However, these methods can be expensive, in terms of both reagents and labor costs as well as time. In recent years, due in part to steady gains in computing power, and especially with the advent of GPU-based calculations, computational methods for predicting the effects of mutations on binding affinity and other physical properties have grown in popularity. At the same time, methodological improvements have resulted in improved accuracy across a wide range of methods, including empirical, machine learning-based, and physics-based methods[10, 11, 12, 13, 14, 15, 16, 17, 18, 19, 20].

Free energy perturbation (FEP), first introduced by Zwanzig in 1954 [21], is a physics-based method for calculating changes in free energy between closely related molecular systems using a series of molecular dynamics (MD) simulations. Modern implementations of FEP have been developed to incorporate substantial improvements in throughput and sampling efficiency [22, 23, 24], as well as the accuracy of molecular mechanics force fields used to calculate interatomic forces and system energy [25, 26, 27, 28, 29].

One such modern implementation which is widely used in structure-based drug design is FEP+ [30]. Over the past decade FEP+ has demonstrated a high level of accuracy and reliability in predicting the binding potencies of small molecules with their protein targets, both via relative binding FEP for congeneric ligands in lead optimization and via absolute binding FEP for diverse ligands in hit discovery [31, 32]. Such calculations have generated positive impact on a large number of prospective studies in active drug discovery projects, leading to faster project progression through faster identification of novel potent chemical matter [33].

Initially developed for ligand perturbations in a protein receptor binding site, the FEP+ method has been adapted to a variety of different applications. We have previously published proof-of-concept studies for the application of FEP+ to predict the effects of neutral and charge-changing perturbations on protein-protein binding affinity [17, 18], as well as protein thermostability calculations to predict melting temperatures (Tm) for mutant proteins [34, 35, 36]. FEP+ has been demonstrated to calculate both protein residue pKa values and free energies between different receptor conformations with high accuracy [37, 38]. It has also been used recently to probe the effects of mutations on the interaction between the severe acute respiratory syndrome coronavirus 2 (SARS-CoV-2) viral spike protein and the human angiotensin converting enzyme 2 (ACE2) receptor, where it outperformed several other computational methods, as determined both by the agreement of its results with experiment and by its ability to identify stabilizing mutations [39]; and to elucidate the structural basis for the selectivity of α-conotoxin variants for different nicotinic acetylcholine receptor subtypes [40].

The aim of the current study is to benchmark the accuracy of FEP+ for protein-protein binding affinity, identify limitations of the current method that need further improvement, and demonstrate how the calculations might be utilized during a protein interface optimization project. We test FEP+ on a curated benchmark data set of binding free energies for single mutational variants of protein-protein binding systems collected from public sources and supplement it with several new experimental measurements from our own work. We apply default FEP+ protocols to assess the accuracy and reliability of the “out-of-the-box” calculations, and systematically investigate cases with large errors. We present an automated protocol for detecting probable outlier cases, to guide the user to more carefully examine the results of cases that satisfy certain chemical and structural criteria in prospective studies. (In this manuscript we use the term “outlier” to refer to cases with absolute errors in relative binding free energy, ΔΔ*G*, larger than 2 kcal/mol relative to experiment.) For one class of outliers, consisting of cases which result in an unpaired buried charge, we introduce a single-parameter empirical correction to account for incomplete relaxation of those systems. Finally, we apply this automated protocol to data from three formerly active biologics design projects to demonstrate the potential value of the method in real-world interface optimization scenarios.

## Results

### Benchmark dataset curation

To assess the performance of protein FEP+ on a range of systems and protein types, we assembled a benchmark dataset of binding affinity measurements from publicly available sources. For each system, we included binding affinity data for the wild-type (*wt*) protein complex as well as for the complex with various single amino acid residue mutations. In total we collected binding data for 9 systems. Binding data for 6 systems were obtained from the SKEMPI 2.0 database [41], along with 3 systems from our own previous work [42, 43, 39], including several previously unpublished measurements (Supplemental Figure 1). To increase the reliability of the experimental values, we limited the dataset to include only measurements made by isothermal calorimetry (ITC) or surface plasmon resonance (SPR). The selected benchmark systems and a summary of the included mutations are presented in Figure 1.

**Figure 1:**
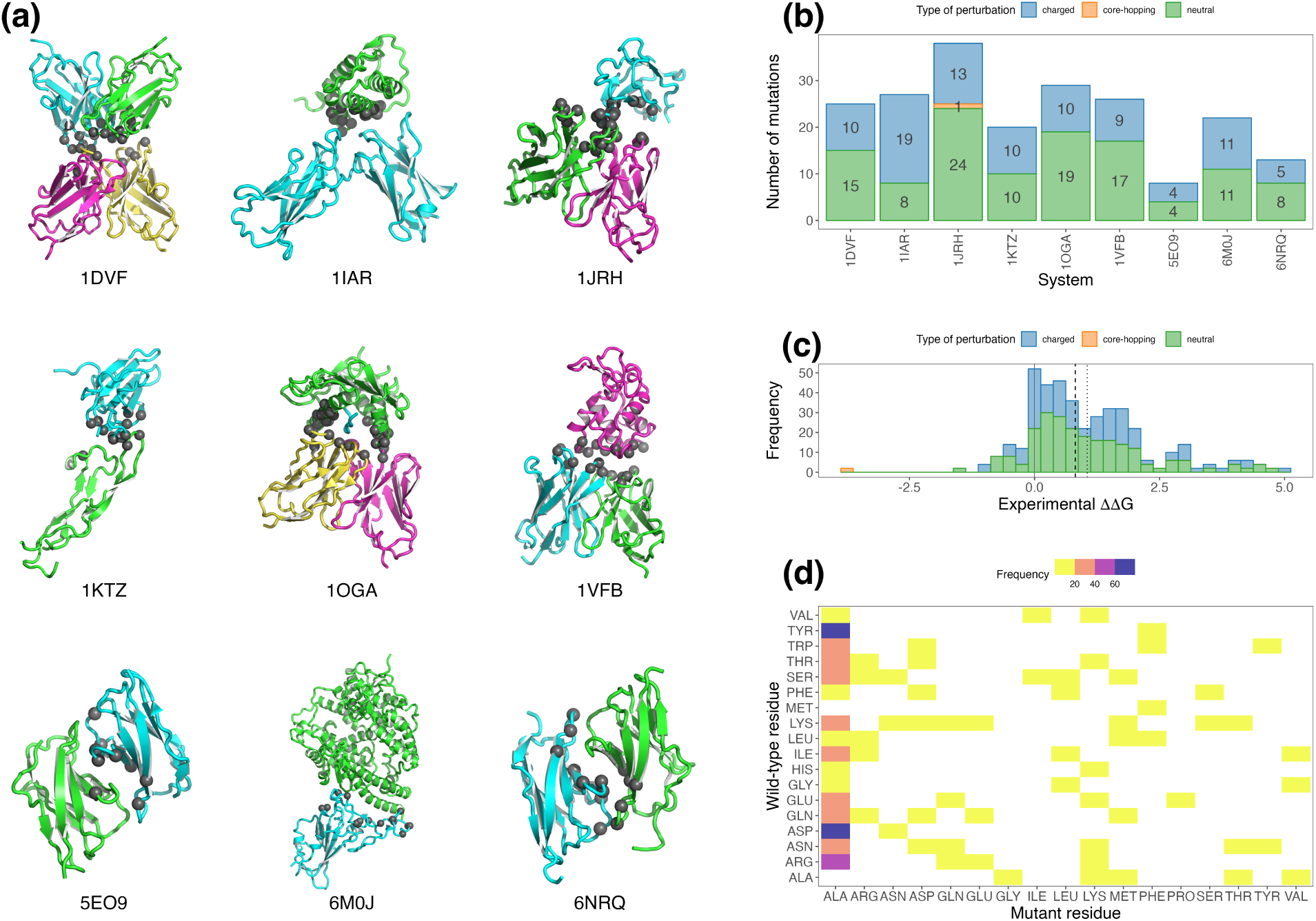
Overview of systems and mutations in the benchmark dataset. (a) Ribbon representation of each model in the benchmark dataset, colored by chain, with Cα positions of mutated residues shown as dark grey spheres. Original PDB accession codes are indicated. (b) Number of mutations per system, colored by FEP perturbation type. (c) Distribution of experimental binding ΔΔ*G* values across all systems, also colored by perturbation type. Median and mean experimental values are shown as dashed and dotted vertical lines, respectively. (d) Heat map showing coverage of amino acid mutation space. Due to the distribution of mutations in the underlying experimental data, mutations to ALA are visibly over-represented.

The full benchmark dataset comprises 208 cases, with 116 neutral perturbation cases, which involve FEP mutation edges between neutral, non-titratable residues; 91 charged perturbation cases, which involve one or more residues with nonzero formal charge (for the current study, these are anionic forms of Asp and Glu; and the cationic forms of His, Lys, and Arg); and 1 “core-hopping” perturbation case, which involves proline and the corresponding change in protein backbone topology. The range of binding ΔΔ*G* values relative to *wt* for the full dataset is −3.79 to +4.91 kcal/mol, with mean and median mutational binding ΔΔ*G* values of +1.05 and +0.81 kcal/mol, respectively. Per-system statistics are presented in Table 1.

**Table 1:** Summary of experimental data for each system in the benchmark dataset. Binding ΔΔ*G* values are given in kcal/mol.

| System | Description | Binding $\Delta\Delta G$ | | | | Number of mutations | | | |
| --- | --- | --- | --- | --- | --- | --- | --- | --- | --- |
|  |  | Min | Mean | Max | Range | Neutral | Charged | Core-hopping | Total |
| 1DVF | IgG1-κ D1.3 Fv / E5.2 Fv | 0.34 | 2.15 | 4.74 | 4.40 | 15 | 10 | 0 | 25 |
| 1IAR | Interleukin-4 / Interleukin-4 receptor | -0.58 | 0.88 | 3.75 | 4.33 | 8 | 19 | 0 | 27 |
| 1JRH | mAb A6 / Interferon-γ receptor | -3.79 | 1.39 | 4.91 | 8.70 | 24 | 13 | 1 | 38 |
| 1KTZ | TGF-β3 / TGF-β type II receptor | 0.22 | 1.56 | 4.04 | 3.82 | 10 | 10 | 0 | 20 |
| 1OGA | HLA-A2 + peptide / JM22 T-cell receptor | -0.46 | 0.53 | 1.93 | 2.39 | 19 | 10 | 0 | 29 |
| 1VFB | IgG1-κ D1.3 Fv / HEW Lysozyme | -0.21 | 0.95 | 3.04 | 3.25 | 17 | 9 | 0 | 26 |
| 5EO9 | Dpr6 / DIP-α | -0.91 | 0.75 | 2.15 | 3.06 | 4 | 4 | 0 | 8 |
| 6M0J | SARS-CoV-2 RBD / human ACE2 | -0.75 | 0.05 | 0.62 | 1.37 | 11 | 11 | 0 | 22 |
| 6NRQ | Dpr10/DIP-α | -1.39 | 0.80 | 2.18 | 3.57 | 8 | 5 | 0 | 13 |
| All |  | -3.79 | 1.05 | 4.91 | 8.70 | 116 | 91 | 1 | 208 |

### Retrospective FEP results

We ran retrospective protein FEP+ calculations for the systems in the benchmark dataset using all-atom structural models derived from the structures deposited with the RCSB Protein Data Bank (PDB), and prepared as described in the Methods section, including the addition of hydrogen atoms and assignment of protonation states expected to be dominant in the bound complex at the pH of the experimental measurements. For each system, a perturbation map was constructed as previously described [44], as a network graph with nodes representing unique variants and edges representing FEP+ perturbations (mutations) between node endpoints. We included perturbations from the starting model to all alternate protonation states of the perturbed residue for mutations to or from titratable amino acids Asp, Glu, His, and Lys. To investigate the convergence of the calculations, the FEP+ simulations were run for 100 ns, then post-processed to obtain the ΔΔ*G* values for each perturbation edge at 10 ns. This post-processing was functionally equivalent to running initial 10 ns simulations followed by extension of each simulation to 100 ns.

The inclusion of alternate protonation states afforded us several options for analyzing binding ΔΔ*G* values for mutations involving titratable residues. In an initial, simplistic treatment of these protonation states, we considered only the initial starting protonation state present in the prepared all-atom structural model, ignoring other possible protonation states for titratable *wt* residues, and reported the calculated ΔΔ*G* value for the perturbation to a predefined protonation state of the mutant residue. Specifically, for this “naïve” treatment, for all mutations to the titratable residues indicated above, we used the binding ΔΔ*G* values for mutations to negatively charged Asp (three-letter residue code `ASP`) and Glu (`GLU`); positively charged Lys (`LYS`); and the neutral tautomer of His that is protonated at the 𝜀 nitrogen (`HIE`). Binding ΔΔ*G* values to other protonation states (neutral `ASH`, `GLH`, `LYN`, and `HID`; and positively charged `HIP`), and the states of titratable *wt* residues not represented by the prepared input model were not considered.

Results for 10 and 100 ns FEP+ calculations with naïve treatment of protonation states are shown in the upper panels of Figure 2. Overall root mean square error (RMSE) for these naïve calculated ΔΔ*G* values compared to experimentally measured values are 1.65 and 1.42 kcal/mol, and coefficient of determination (R^2^) values are 0.29 and 0.27, for 10 and 100 ns results, respectively. A report of per-system dataset statistics can be found in Table 2.

**Figure 2:**
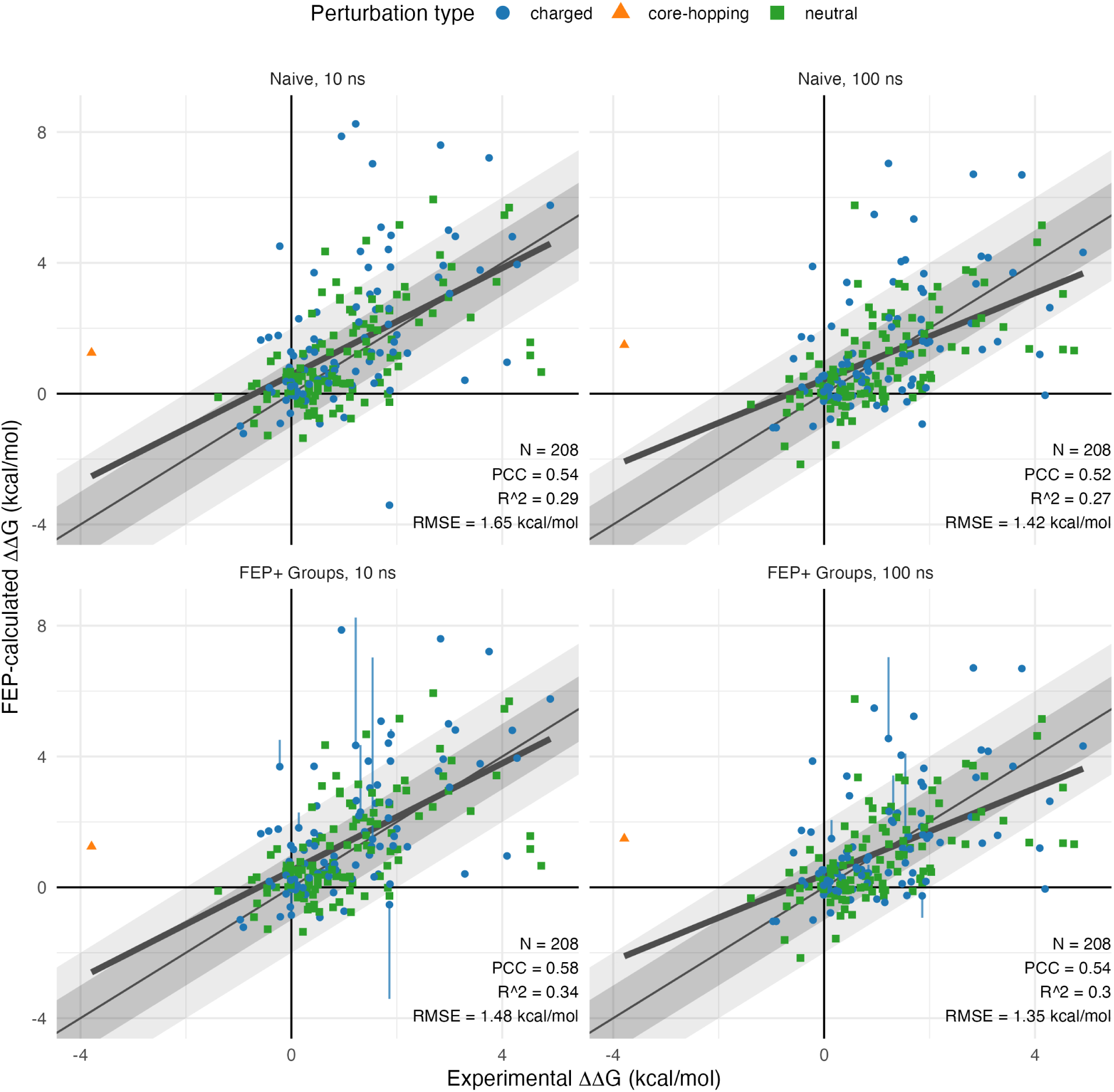
FEP+ results versus experiment. Correlation plots of FEP+ predicted binding ΔΔ*G* vs. experimental binding ΔΔ*G* relative to WT for the benchmark dataset mutations. Results are shown for 10 ns (left) and 100 ns (right) FEP+ calculations, with both naïve (top) and FEP+ Groups (bottom) treatment of protonation states. Grey and light grey diagonal shaded areas represent regions of absolute error less than 1 and 2 kcal/mol, respectively. A least squares best fit line (bold) and relevant correlation statistics are indicated for each plot.

**Table 2:** Summary of retrospective FEP+ results compared to experimental data for each system in the benchmark dataset. Predicted binding ΔΔ*G* error and correlation statistics are shown for 10 and 100 ns simulations, with naïve and FEP+ Groups treatment of protonation states for each. Root mean square error (RMSE) values are given in kcal/mol.

| System | 10 ns |  |  |  | 100 ns |  |  |  |
| --- | --- | --- | --- | --- | --- | --- | --- | --- |
|  | Naïve |  | FEP+ Groups |  | Naïve |  | FEP+ Groups |  |
| | RMSE | $R^2$ | RMSE | $R^2$ | RMSE | $R^2$ | RMSE | $R^2$ |
| 1DVF | 1.83 | 0.30 | 1.57 | 0.36 | 1.74 | 0.26 | 1.70 | 0.27 |
| 1IAR | 2.45 | 0.44 | 1.97 | 0.57 | 2.04 | 0.50 | 1.76 | 0.57 |
| 1JRH | 1.83 | 0.26 | 1.82 | 0.26 | 1.78 | 0.20 | 1.77 | 0.21 |
| 1KTZ | 0.87 | 0.74 | 0.87 | 0.74 | 0.85 | 0.78 | 0.85 | 0.78 |
| 1OGA | 0.61 | 0.15 | 0.63 | 0.19 | 0.68 | 0.08 | 0.68 | 0.12 |
| 1VFB | 1.86 | 0.33 | 1.86 | 0.33 | 1.36 | 0.27 | 1.36 | 0.27 |
| 5EO9 | 1.55 | 0.61 | 1.55 | 0.61 | 1.02 | 0.78 | 1.02 | 0.77 |
| 6M0J | 0.66 | 0.31 | 0.66 | 0.31 | 0.67 | 0.47 | 0.67 | 0.47 |
| 6NRQ | 1.95 | 0.54 | 1.22 | 0.66 | 1.05 | 0.64 | 0.80 | 0.67 |

We found that on average the longer 100 ns simulations resulted in smaller overall errors, but similar correlation coefficients, compared to the more typical 10 ns simulations. This was not surprising, as the cases where initial configurations of the mutants were in high energy conformations were partially relaxed in the 100 ns simulations, but likely required even longer time for full relaxation. Notably, relatively few cases showed substantial change in predicted binding ΔΔ*G* between the 10 ns and 100 ns timepoints, with median absolute ΔΔ*G* shifts of 0.33 kcal/mol, as shown in Supplemental Figure 2.

### Effect of Protein FEP+ Groups treatment

Several of the large outliers among the naïve treatment results involved mutation to or from titratable residues. The preferred protonation states of these titratable residues might differ between the relevant physical macrostates, namely the bound protein-protein complex and the unbound protein in solution. In the physical system, these alternate protonation microstates are able to interconvert freely, with the populations of the protonated or deprotonated microstate of each titratable residue at the experimental pH being determined by its pKa in the given physical macrostate. A more robust approach to analyzing such mutations requires considering all microstates as a single statistical mechanical ensemble, to better reflect the physical system that was tested experimentally. The method we developed to implement this robust treatment of alternate protonation states, called Protein FEP+ Groups, groups together nodes (mutational variants) in the FEP+ perturbation map that represent the same physical amino acid sequence but differ in modeled protonation state, and a single, population-based, Boltzmann-weighted predicted binding ΔΔ*G* is calculated for the entire ensemble of protonation microstates in the node group. The pH-dependent per-site population of each protonation microstate is obtained via the calculated pKa of the specific amino acid residue at the given titratable site (residue position) in the given physical macrostate. The Protein FEP+ Groups treatment is described in more detail in the Methods section below.

Two examples of FEP+ Groups treatment are presented in Figure 3. As shown in Figure 3(a), 1IAR A:W91 was mutated to Asp, so we considered both the deprotonated `ASP` as well as protonated `ASH` forms for the mutant. Experimentally the Asp mutation yielded a binding ΔΔ*G* of +1.31 kcal/mol, but the naïve prediction of +3.42 was overly unfavorable. When all protonation states were considered, the largely hydrophobic surrounding environment resulted in unfavorable binding ΔΔ*G*s for both the charged and neutral forms, but the neutral `ASH` form was favored over `ASP` by nearly 3 kcal/mol in the binding interface, despite the presence of a nearby Arg side chain. FEP+ Groups-calculated pKas of the mutant Asp in the unbound protein and in the context of the protein-protein binding interface indicated a change in preferred protonation state upon binding at the experimental pH of 7.2. In the unbound state, the deprotonated `ASP` form was calculated to be dominant, with an A:D91 side chain pKa of 6.5; whereas in the bound state, a higher complex pKa of 8.5 indicated a preference for the neutral `ASH` form. Accounting for this change in dominant protonation state, the FEP+ Groups treatment yielded a more accurate predicted ΔΔ*G* of +1.99 kcal/mol, corresponding to a decrease in absolute error from 2.11 to 0.68 kcal/mol.

**Figure 3:**
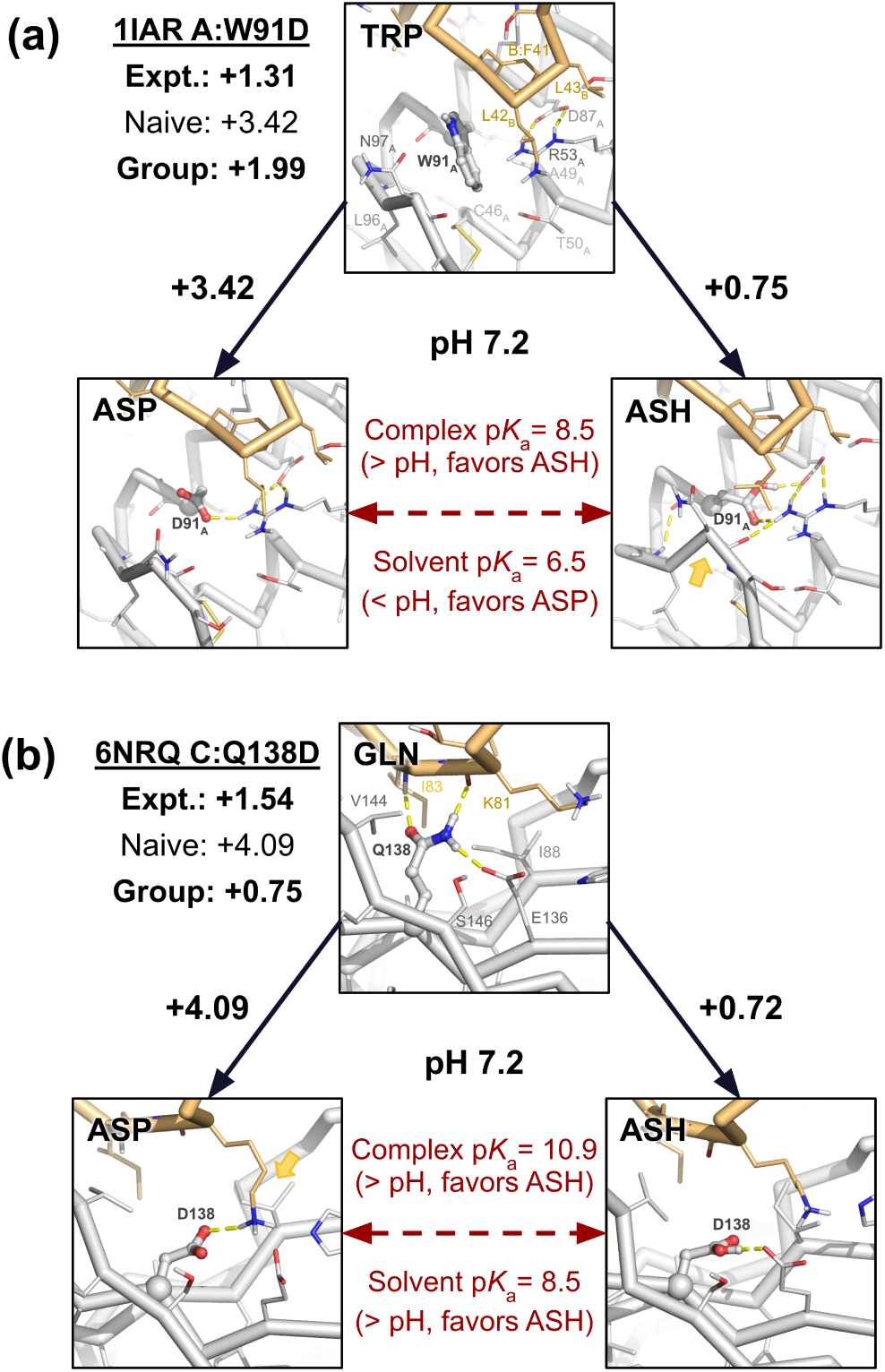
Protein FEP+ Groups treatment example cases. Ribbon representations of either the *wt* input structure or a representative mutant trajectory frame, with the chain(s) for the binding component that includes the mutation shown in light grey, the binding partner chain(s) in tan, and key polar interactions as dashed yellow lines. FEP-calculated 100 ns binding ΔΔ*G* values for mutation edges are shown along solid arrows, and FEP-based calculated pKas for titration edges in the bound (complex) and unbound (solvent) forms are shown along the horizontal dashed arrows. Experimental, naïve, and FEP groups-treated ΔΔ*G* values are listed in kcal/mol. (a) For 1IAR A:W91D the calculated pKas for the mutant Asp91 carboxylate indicated a change in dominant protonation state between the unbound and bound forms at experimental pH. FEP+ Groups treatment resulted in reduced absolute error compared to the naive result. (b) The 6NRQ C:Q138D mutant Asp138 carboxylate had elevated FEP-calculated pKa values in both unbound and bound states, reflecting the largely hydrophobic residue environment, limited solvent access, and the conformational change required to form a salt bridge with D:K81. FEP+ Groups treatment again increased accuracy with respect to the experimental value.

A second example, shown in Figure 3(b), is 6NRQ C:Q138D. This mutation was also unfavorable experimentally (+1.54 kcal/mol), but the naïve predicted value was again too highly destabilizing (+4.09). In the crystal structure, the *wt* C:Gln138 side chain acts both as an H-bond donor (to the C:Glu136 side chain as well as to the D:Lys81 backbone carbonyl) and as an H-bond acceptor (from D:Ile83 backbone N–H). Mutation to either Asp state was predicted to be unfavorable, due to the disruption of these interactions. FEP+ Groups treatment produced a predicted ΔΔ*G* of +0.75, reducing the error from 2.55 to 0.79 kcal/mol. Due to hydrophobic contacts and the proximity of C:Glu136, FEP+ Groups-calculated pKas for both physical macrostates were significantly elevated compared to the pKa value of 3.67 for an isolated Asp in solution [45], with calculated pKa values of 8.4 (unbound) and 10.9 (bound), thus indicating the neutral `ASH` form was preferred in both unbound and bound states at the experimental pH of 7.2. Accordingly, in this case mutating to `ASH` alone would have provided a good estimate of the FEP+ Groups-treated value, due to the minimal contribution of the high-energy (and therefore low-prevalence) `ASP` state, though this would not necessarily have been apparent based on the input structure alone.

After applying the FEP+ Groups treatment to the retrospective calculations, we observed decreased RMSE and increased Pearson correlation coefficient (PCC) and R^2^ at both 10 and 100 ns time points. These results are presented in the lower plots of Figure 2. The number of cases where this groups treatment substantially affects the results is relatively small, with differences larger than 0.25 kcal/mol observed among the 100 mutations involving titratable residues for 8 and 6 cases, for the 10 and 100 ns results, respectively. Importantly, Protein FEP+ Groups treatment yielded calculated ΔΔ*G* values with reduced absolute error compared to experiment on average: among these cases involving titratable residues, FEP+ Groups treatment resulted in decreased RMSE at both timepoints, from 2.33 to 1.48 kcal/mol at 10 ns, and from 1.83 to 1.35 kcal/mol at 100 ns.

### Analysis of outliers

The FEP+ results from longer simulation time and rigorous treatment of titratable residues yielded accuracy consistent with results from previous studies, with RMSE in the range of about 1.0-1.5 kcal/mol [17, 18]. However, a number of mutations exhibited absolute errors in binding ΔΔ*G* larger than 2 kcal/mol compared to experiment, which would reduce the prediction accuracy in prospective studies. In an effort to investigate the limitations of the current default protein binding FEP+ methodology and guide future methods improvement, we systematically examined the cases from the 100 ns FEP+ Groups-treated results with the largest errors relative to experiment and identified common features for these outliers which might be useful for prospective application of the method. We initially focused on cases with absolute errors larger than 2 kcal/mol, which totaled 19 cases, or about 9% of the full dataset. Close inspection of the *wt* crystal structures, mutant FEP+ starting models, and FEP+ trajectories for these cases suggested several distinct sources of error, which are summarized in Table 3. These error sources can broadly be categorized as either sampling errors, where the system had not yet fully relaxed or failed to sample relevant areas of conformation space; or force field errors, where specific interaction types may not be handled properly by the current OPLS4 force field. Examples of the various outlier classes are shown in Figure 4.

**Figure 4:**
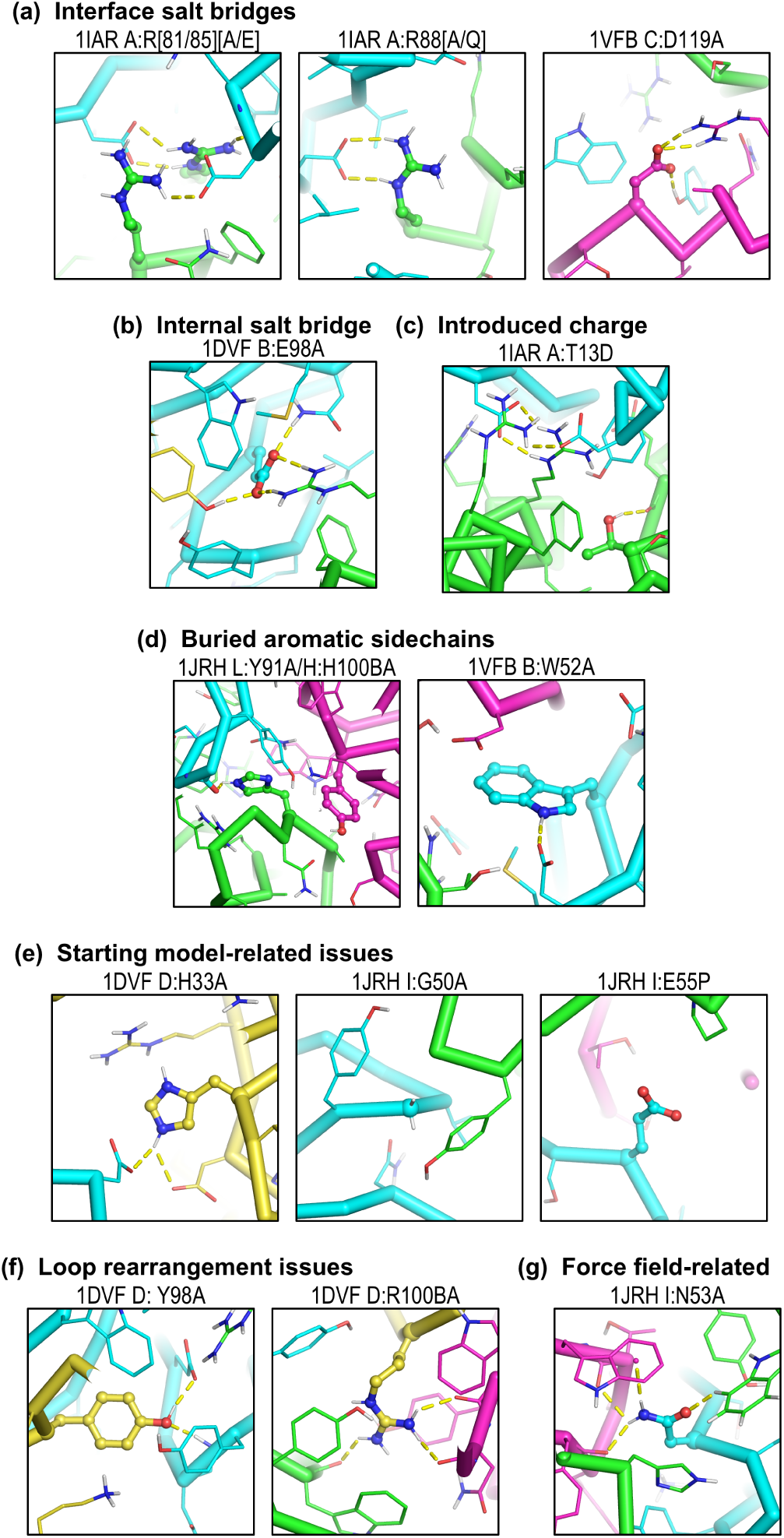
Classes of outlier cases with absolute errors greater than 2 kcal/mol. Structural images in ribbon representation, with key *wt* sidechains shown as thin sticks, colored by chain and atom type. Mutations are indicated above each panel and mutated residues are shown in ball-and-stick representation. (a) Salt bridge-breaking cases include *trans* (left two panels) and *cis* (second from right panel) interface salt bridges, and an internal salt bridge between the heavy and light chains of an antibody Fv region (right panel). (b) Two outlier cases introduced a buried charge. (c) A variety of sampling related cases. (d) Outliers resulting from mutation of buried aromatic residues to much smaller alanine. (e) One force field-related case involved a disrupted π-hydrogen bond interaction.

**Table 3:** Manual classification of outlier cases with absolute error > 2 kcal/mol relative to experiment. Cases within each outlier class are sorted by decreasing absolute error. All ΔΔ*G* and error values are given in kcal/mol.

| Error type | Outlier class | System | Mutation | Exp. $\Delta\Delta G$ | Pred. $\Delta\Delta G$ | Error | Abs. Error |
| --- | --- | --- | --- | --- | --- | --- | --- |
| Sampling | Disrupted salt bridge (interfacial, *trans*) | 1IAR | A:R88Q | 2.83 | 6.71 | 3.88 | 3.88 |
|  |  | 1IAR | A:R85E | 1.22 | 4.55 | 3.33 | 3.33 |
|  |  | 1IAR | A:R85A | 0.43 | 3.40 | 2.97 | 2.97 |
|  |  | 1IAR | A:R88A | 3.75 | 6.69 | 2.94 | 2.94 |
|  |  | 1IAR | A:R81E | 1.46 | 4.04 | 2.58 | 2.58 |
|  |  | 1IAR | A:R81A | 0.48 | 2.80 | 2.32 | 2.32 |
|  | Disrupted salt bridge (interfacial, *cis*) | 1VFB | C:D119A | 0.95 | 5.48 | 4.53 | 4.53 |
|  | Disrupted salt bridge (internal) | 1DVF | B:E98A | 4.19 | -0.05 | -4.24 | 4.24 |
|  | Introduced charge | 1IAR | A:T13D | -0.22 | 3.86 | 4.08 | 4.08 |
|  | Buried aromatic, large size change | 1JRH | L:Y91A | 0.58 | 5.76 | 5.18 | 5.18 |
|  |  | 1JRH | H:H100BA | 1.70 | 5.23 | 3.53 | 3.53 |
|  |  | 1VFB | B:W52A | 0.64 | 3.36 | 2.72 | 2.72 |
|  | Problematic starting conformation | 1JRH | I:G50A | 4.53 | 1.35 | -3.18 | 3.18 |
|  |  | 1JRH | I:E55A | -0.43 | 1.74 | 2.17 | 2.17 |
|  |  | 1DVF | D:H33A | 1.86 | -0.26 | -2.12 | 2.12 |
|  | Loop rearrangement | 1DVF | D:Y98A | 4.74 | 1.32 | -3.42 | 3.42 |
|  |  | 1DVF | D:R100BA | 4.09 | 1.20 | -2.89 | 2.89 |
|  | Proline | 1JRH | I:E55P | -3.79 | 1.49 | 5.28 | 5.28 |
| Force Field | Disrupted $\pi$ interaction | 1JRH | I:N53A | 3.89 | 1.37 | -2.52 | 2.52 |

### Outliers due to insufficient sampling and other conformational issues

The largest outlier class comprised 7 cases, one-third of the outlier cases, where mutation resulted in the elimination of a salt bridge present at the binding interface in the *wt* structure. These are depicted in Figure 4(a). All seven cases were unfavorable experimentally, with binding ΔΔ*G*s ranging from +0.43 to +3.75 kcal/mol, but FEP+ consistently predicted them to be too unfavorable with signed errors ranging from +2.3 to +4.5 kcal/mol. Six of the cases (which were all mutations at one of three Arg positions in the 1IAR system: A:R81A/E, A:R85A/E, and A:R88A/Q) disrupted *trans* salt bridges, that is, salt bridges involving residues from both protein binding components, whereas one case (1VFB C:D119A) disrupted a *cis* salt bridge, between two residues from the same side of the interface. We observed that the mutated residues in these outlier cases were largely buried, such that upon mutation of one of the two residues forming a salt bridge, the remaining partner residue was left in an unstable configuration: a charged residue, buried at the interface with limited solvent accessibility. FEP+ correctly predicted unfavorable positive binding ΔΔ*G* values for all 7 of these cases, but dramatically overestimated the magnitude: for the 6 *trans* salt bridge cases, FEP+ overpredicted the unfavorability of the mutations by approximately 3 kcal/mol.

Two related outlier classes also involved unstable charged configurations upon mutation. The first of these, shown in Figure 4(b), involved a single case where a salt bridge was disrupted upon mutation, but the mutated residue was entirely buried within the mutated protein. Specifically, for 1DVF B:E98A (expt. +4.19; pred. −0.05), in the *wt* structure residue B:Glu98 forms a salt bridge with A:Arg96 across the internal V_H_-V_L_ heterodimerization interface, between the heavy and light chain variable domains of the antibody Fv region. Although close to the binding interface, this residue was buried in the unbound antibody and not in contact with the binding partner, and therefore not directly involved in binding. The mutation disrupting the salt bridge between the heavy and light chains of the antibody could significantly change the relative orientation between the V_H_ and V_L_ domains, thus disrupting the binding with binding partner. However, the relatively short timescale (100 ns) in FEP+ simulations was not sufficient to fully relax the conformations of and relative orientation between the mutant V_H_ and V_L_ domains in the unbound mutant structure, resulting in an underprediction of the unfavorability of the mutation and a large absolute error of 4.24 kcal/mol. The final charge-related outlier, shown in Figure 4(c) was 1IAR A:T13D (expt. -0.22; pred. +3.86) where a neutral residue was mutated to a charged amino acid at a largely buried position at the protein-protein complex interface. This sort of charge-introducing mutation is functionally similar to the salt bridge-breaking cases, in that the end result is an unpaired charge in an unfavorable configuration, which likely requires substantial relaxation to adopt a conformation not accessible on typical FEP+ timescales. We noted that there were also several charge-flipping cases in our dataset, whose mutations inverted the sign of the charge on the amino acid side chain, resulting in a net change in formal charge of ±2. Two of these (1IAR A:R85E and A:R81E) were among the salt bridge-breaking cases described above, but the other two (1IAR A:K12E and A:K84D), were also unfavorable experimentally and gave signed errors greater than +1 kcal/mol, suggesting that these types of buried (or mostly buried) unpaired charges resulting from charge flipping perturbations also tend to be undersampled.

Figure 4(d) depicts an outlier class conisiting of three cases (1JRH L:Y91A and H:H100BA; and 1VFB B:W52A) which involved mutations of larger aromatic residues, buried in the complex structure, to much smaller side chains, namely alanine. All three were modestly unfavorable experimentally (+0.6 to +1.7 kcal/mol), but FEP+ overpredicted the unfavorability by 3-5 kcal/mol. Due to their large surface area and low solvent accessibility, the aromatic residue side chains at these positions made numerous interactions in their respective *wt* structures, including hydrophobic interactions, hydrogen bonds, and π interactions. When such large side chains were eliminated from the structure entirely (rather than being replaced with a similarly sized amino acid), the 100 ns FEP+ simulations appeared not to have been sufficiently long to allow for full relaxation and rearrangement of the surrounding residues into the resulting void, thereby producing FEP+ predictions that were too highly unfavorable. This type of slow convergence behavior was previously observed for `TRP->ALA` mutations in a dataset with anti-HIV-1 antibodies in complex with HIV-1 gp120 spike proteins [17]. Notably, the *wt* residues for the two 1JRH cases here (L:Tyr91 and H:His100B) are adjacent and interact in the complex, suggesting a common mode of error for both.

An outlier class with errors derived from sampling issues related to starting model conformations, shown in Figure 4(e), included 1DVF D:H33A and three mutations on a loop in the 1JRH system, namely I:G50A, I:E55A, and I:E55P. In the 1DVF D:H33A case (expt. +1.86, pred. -0.26), which is presented in more detail in Supplemental Figure 3, the geometry of the *wt* D:His55 residue (modeled as `HIP`) from the crystal structure showed its N𝜀 proton interacting with two side chain carboxylate groups, from D:Asp52 (in *cis*) and B:Asp100 (in *trans*). In the 100 ns FEP+ simulation, this initial geometry resulted in poorly defined, low-persistence interactions with the two carboxylate groups for approximately the first 50 ns of the simulation, after which a stable configuration appeared with the His55 ring flipped 180° and its N𝛿 and N𝜀 protons each forming hydrogen-bonding salt bridge interactions with one of the two Asp side chains. The lack of these interactions for much of the the *wt* trajectory from the FEP+ calculations resulted in an underestimation of the unfavorability of the mutation. A subsequent standard MD simulation of the *wt* structure resulted in rapid adoption of the stable, dual-salt bridge configuration of the imidazole ring, which remained stable throughout the trajectory. Furthermore, running the D:H33A mutation by FEP+ using the final frame of this MD trajectory as the input model yielded an accurate binding ΔΔ*G* prediction of +1.56 kcal/mol, within 0.30 kcal/mol of the experimentally determined value of +1.86 kcal/mol. This result suggests the starting conformation of D:His33 may be the result of a modeling error in the original crystal structure; unfortunately the 1DVF PDB entry has no associated experimental diffraction data, so it is not possible to ascertain the degree to which this FEP+ and MD-validated conformation of D:His33 might be more consistent with the experimental crystal context than the originally deposited conformation.

The 1JRH I:G50A (expt. +4.53, pred. +1.49), I:E55A (expt. -0.43, pred. +1.74), and I:E55P (expt. -3.79, pred +1.63) cases, which are also depicted in Supplemental Figure 4, each involve mutation of a residue on the flexible I:50-56 loop of the interferon γ receptor 1 (IFNγR1) antigen. The first of these, 1JRH I:G50, is located in a tight turn with geometry resembling a γ-turn, and backbone φ and ψ angles that make it an outlier in the Ramachandran plot for the crystal structure (Supplemental Figure 4 a-b) This high-energy backbone conformation would require substantial relaxation to achieve the true lowest energy conformation. Indeed, two other crystal structures (1FYH chain B, and 6E3L chain D) show IFNγR1 adopting an alternative conformation of the I:50-56 loop, further suggesting that the conformation in the 1JRH crystal structure may be suboptimal. The second mutation site on this loop, 1JRH I:E55, has a side chain that is highly solvent exposed in the complex, making no direct contact with any antibody residue in the *wt* crystal structure. It is approximately 6 Å from two different primary amino groups—namely the solvent-exposed (and therefore flexible) H:Lys64 side chain and the N-terminal amino group from L:Ser1—and FEP+ and MD trajectories show it interacting with the charged N-terminus of chain L. Mutation to Ala eliminated this charged interaction, thus accounting for the unfavorable prediction by FEP+. In this case, where the starting structure is substantially unrelaxed, random variation in the direction and degree of relaxation among different simulations can lead to high levels of noise in predicted results as compared to experiment. In other words, the interaction with the charged chain L N-terminus may not be representative of the physical system. More generally, for mutation of any non-contact residue in a crystal structure to yield a binding ΔΔ*G* significantly different from zero essentially requires some conformational rearrangement, either in the unbound form, the bound form, or both. Accurate simulation of such conformational changes would necessitate sampling beyond what is achievable in the standard protein binding FEP+ workflow, thus rendering cases of this type not typically suitable for protein binding FEP+ simulation using standard protocols. Notably mutations of 1JRH I:G50 to Ala; and I:E55 to Ala and Pro result in errors of similarly large magnitude, but opposite signs. Whereas we attribute the negative signed error of the I:G50A mutation to initial strain in the *wt* complex; the large positive signed errors for the I:E55 mutations to Ala and especially Pro in this flexible loop suggest that further relaxation of the mutant bound complex is required, and is more pronounced in the Pro case, likely due to a change in the loop backbone conformation required to compensate for the rigidity of the Pro backbone.

A final sampling-related outlier class, shown in Figure 4(f), involved mutations in the 1DVF antibody CDR H3 loop, D:Y98A (expt. +4.74, pred. +1.32) and D:R100BY (expt. +4.09, pred. +1.2). These outliers exhibited the negative signed FEP error common to cases involving unsampled rearrangement in the unbound mutant protein. Both cases were predicted to be modestly unfavorable, but experimentally were much more unfavorable. For each of these cases, we repredicted the conformation of the CDR H3 loop in the unbound free antibody, and in each case identified a different loop conformation with lower energy than the crystal structure-like conformation (Supplemental Figure 5). These results suggest the error in the FEP+ predictions may be at least partially due to this type of unsampled H3 loop rearrangement in the unbound mutant protein.

### Outlier attributed to force field limitations

For one outlier case, we hypothesized that the deviation from the experimentally measured binding ΔΔ*G* value resulted from a limitation of the fixed-charge force field used in FEP+ simulations. More specifically, the case in question, 1JRH I:N53A (expt. +3.89, pred. +1.37), involved mutation of a residue that participates in a well-defined π interaction in the *wt* crystal structure. As depicted in Figure 4(g), the π interaction in question, rather than a more typically discussed π-π or π-cation interaction, was instead a π-hydrogen bond [46] between the *wt* Asn sidechain amide and the π electron cloud of the 6-membered aromatic ring of I:Trp96. Due to the fixed-charge nature of the OPLS4 force field, it was not possible for the FEP+ calculation to account for the contribution of this π interaction to the stability of the *wt* complex, which resulted in a predicted binding ΔΔ*G* value that was too favorable by 2.5 kcal/mol.

We investigated this putative force field-related outlier via a 100 ns FEP+ calculations run with a developmental version of the OPLS force field which includes explicit polarization of aromatic residue sidechains. The FEP+ results with explicit polarization resulted in substantial improvement of the predicted binding ΔΔ*G*, to +3.09 kcal/mol (an error of -0.8 compared to experiment). This preliminary result reflects an increased stabilization of the *wt* complex by more accurately modeling the energetics of the π-hydrogen bond between the Asn and Trp sidechains, and suggests the inclusion of explicit polarization in future versions of the OPLS force field may yield improved results in similar cases.

### Automated detection of probable outliers

Based on the above observations about the largest outliers in the benchmark dataset, we developed an automated method for detecting cases where a protein binding FEP+ calculation is likely not able to accurately predict the effect of mutations on the binding affinity. These probable outliers are identified based on information available in the output perturbation map (`.fmp`) file from the protein binding FEP+ job. This map file contains structural information for the starting models used in simulation for both *wt* and mutant nodes (protein variants), as well as free energy time series data and a summary report of interactions made by the *wt* and mutant residues at the mutated position for each perturbation edge (mutation).

The automated script uses two sets of empirical rules to flag probable outlier cases. The first set identifies cases that break salt bridges or otherwise introduce or flip charges at buried or mostly buried sites. A second set of rules identifies cases that involve either mutation of large aromatic residues to substantially smaller residues at buried sites; or mutation to Proline, which has unique backbone torsional characteristics among the amino acids. These rules use the features derived from the perturbation map as well as a set of five orthogonal, tunable parameters: 1) a fractional solvent-accessible surface area (fSASA) cutoff to classify buried residues; 2) a number of charged neighbors cutoff to identify interacting clusters of charged or titratable amino acids; 3) a residue size change cutoff to identify large changes in the number of heavy (non-hydrogen) atoms; 4) an interaction frequency cutoff as a threshold for defining trajectory-derived interactions; and 5) a predicted ΔΔ*G* cutoff to identify highly destabilizing perturbations. For the results described in this manuscript, we used an fSASA cutoff of 0.1; charged neighbors cutoff of 6; residue size change cutoff of 5; interaction frequency cutoff of 0.5; and ΔΔ*G* cutoff of 2.5 kcal/mol.

The automated outlier prediction script flagged 22 potentially problematic cases among a total of 208 mutations from the 100 ns FEP+ Groups-treated benchmark dataset. This included 12 out of 19 outlier cases with absolute errors > 2 kcal/mol (63%). Additionally we marked as probable outliers 7 cases out of 55 with absolute errors between 1-2 kcal/mol (13%). Of the 134 well predicted cases with absolute errors < 1 kcal/mol, only 3 (2%) were flagged as likely outliers. A summary of the flagged cases is presented in table Table 4.

**Table 4:** Summary of automatic flagging of probable outlier cases by magnitude of FEP+ predicted binding ΔΔ*G* error vs. experiment.

| Error range | Flagged | Unflagged | % Flagged |
| --- | --- | --- | --- |
| >2 kcal/mol | 12 | 7 | 63 |
| 1-2 kcal/mol | 7 | 48 | 13 |
| 0-1 kcal/mol | 3 | 131 | 2 |

### Empirical correction of charged outlier cases

We observed that the magnitudes of the errors for the automatically flagged charge-related outlier cases from the benchmark dataset were relatively consistent across the individual outlier categories, with a mean absolute error of 2.8 ± 1.0 kcal/mol. Indeed, when one considers that the source of the error in each of the categories is the same—namely, insufficient sampling after burial of a charge without an oppositely charged partner residue—this is not unexpected. Therefore, we sought to define an empirical correction term that could be applied generally to such automatically flagged charge-related cases.

In each case, the sign of the error relative to experiment depended only on the location of the offending charged residue, regardless of whether the charge was introduced with the mutation or left unpaired by mutation of a neighboring residue. Specifically, unpaired charges at the binding interface always exhibited a positive signed error in the FEP+ prediction relative to experiment, that is, the predicted binding ΔΔ*G* values were too unfavorable. Whereas, leaving an unpaired charge internal to one of the proteins and not involved in binding, as described above for 1DVF B:E98A, resulted in a negative signed error. This was consistent with our interpretation that in the case of a positive signed error, the mutant bound complex was insufficiently relaxed in simulation, thereby causing the predicted ΔΔ*G* to be too unfavorable. Conversely, for 1DVF B:E98A (which was the only example among these cases with a negative signed error), the antibody Fv must retain or adopt the binding-competent conformation observed in the crystal structure in order to bind the antigen, but in order to alleviate the buried charge at the V_H_-V_L_ interface, the unbound antibody Fv region in the physical system would readily relax to a lower-energy conformation, which was unable to be sampled on the timescale of these calculations. The associated reorganization penalty in the unbound mutant state was missing from the FEP+ calculation, resulting in a predicted ΔΔ*G* value that was too favorable compared to experiment. Given these well-defined criteria for determining the sign of the charged outlier correction term, starting with the predicted ΔΔ*G* value for each case we either subtracted (all binding interface charged outlier cases) or added (1DVF B:E98A only) 2.8 kcal/mol to produce empirically corrected predicted ΔΔ*G* values.

After applying the empirical correction to the charged outlier cases, and excluding the other automatically flagged cases, which we treat as lower confidence predictions more likely to require extensive investigation or non-default FEP+ protocols, or in some cases to be beyond the domain of applicability of FEP+, we observed improved overall error and correlation statistics for the benchmark dataset. These results are presented in Figure 5(a), and demonstrated a marked improvement in both RMSE (which decreased from 1.35 to 1.03) and R^2^ (which increased from 0.3 to 0.42) compared to the full 100 ns FEP+ Groups treated dataset as shown in the lower right panel of Figure 2.

**Figure 5:**
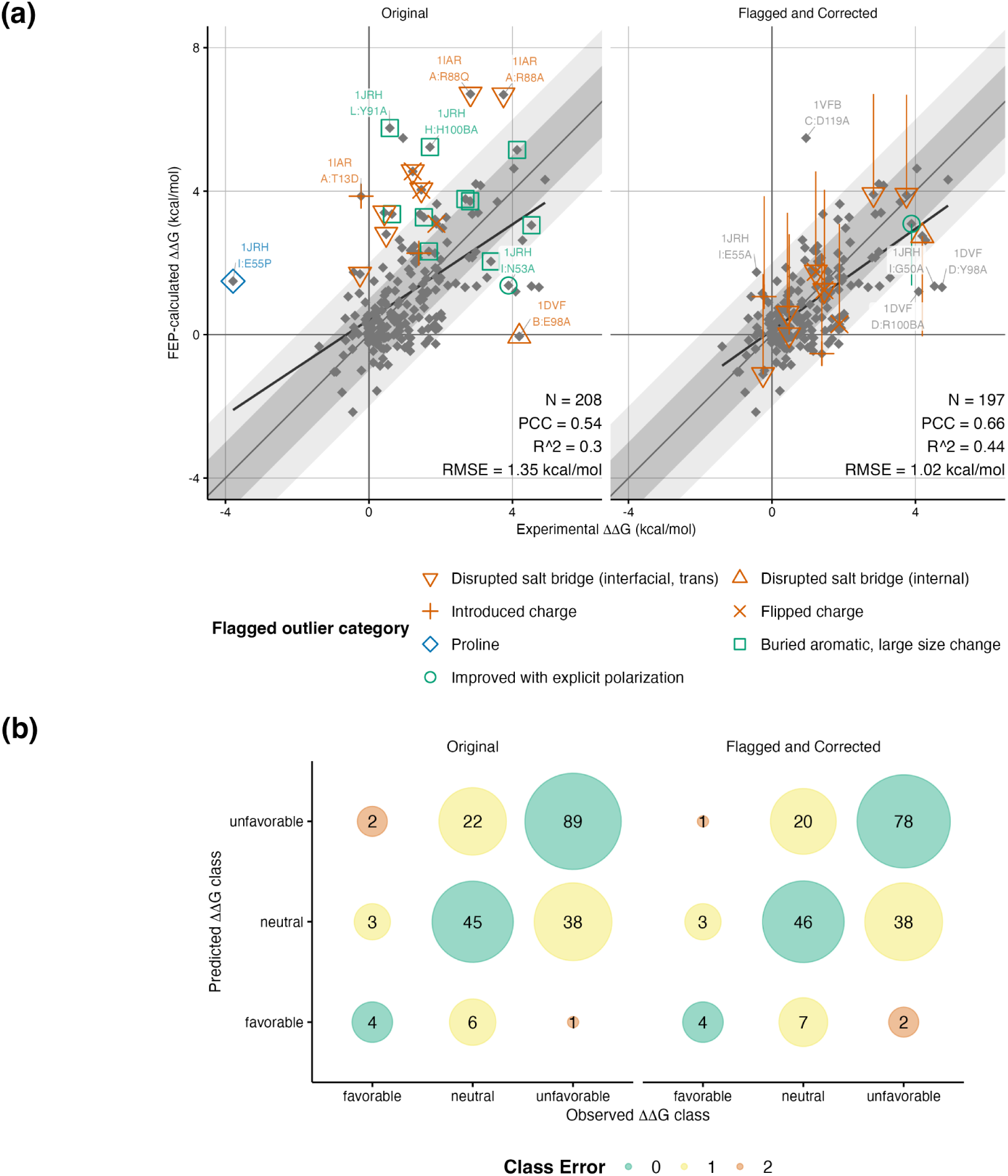
Automated classification and empirical correction of FEP+ outliers for the benchmark dataset. (a) Correlation plots 100 ns FEP+ Groups-treated results vs. experiment, before and after automated outlier flagging and empirical correction of charged outlier cases. Flagged outlier categories are indicated. The direction and magnitude of the applied correction for charged outlier cases is indicated by orange vertical tails. One case (1JRH I:N53A) which demonstrated substantial improvement in preliminary testing of a force field with explicit polarization is indicated by a green circle and a green vertical tail in the right plot. Diagonal shaded areas, least squares fit line, and statistics are shown as in Figure 2. (b) Three-class confusion matrices for the data from (a), with ΔΔG values classified as either favorable (ΔΔG ≤ -0.5 kcal/mol), unfavorable (ΔΔG ≥ 0.5), or neutral (-0.5 < ΔΔG < 0.5).

### Utility of FEP+ calculations for prospective design

For *in silico* free energy predictions to be generally useful in prospective design, they should be able to classify the relative binding ΔΔ*G* values of mutations as favorable, unfavorable, or neutral with good accuracy. In Figure 5(b) we show a 3-class confusion matrix for the 100 ns FEP+ Groups-treated results after applying the empirical correction for charged outlier cases described above and excluding the other automatically flagged cases indicated in Figure 5(a). Mutations were classified as either favorable (binding ΔΔ*G* ≤ -0.5 kcal/mol), neutral (ΔΔ*G* between -0.5 and 0.5 kcal/mol), or unfavorable (ΔΔ*G* ≥ 0.5 kcal/mol), using experimentally observed and FEP-predicted relative binding free energy changes. We found that the FEP+ predictions successfully classified cases from the benchmark dataset with an overall balanced accuracy (arithmetic mean of sensitivity and specificity) of 0.69. Additionally, only 3 outlier cases remained with a classification error larger than 1 class, indicating that the automatic flagging procedure successfully identified for removal from the high confidence dataset many of the predictions with the largest errors.

Computational results are often used as a “pre-filter” step in the molecular design process, to narrow the scope and cost of a project by limiting the number of designs taken forward into the experimental validation stage of the process. For the typical goal of identifying experimentally favorable “active” mutations, if we were to take the full benchmark set as a hypothetical example and consider the most restrictive application of these classification results with only cases having calculated ΔΔ*G* of -0.5 kcal/mol or less (using the unflagged 100 ns cases after FEP+ Groups treatment of protonation states and automated outlier flagging and empirical correction) to be tested by experiment, this would result in the experimental testing of only 13 variants, with a positive hit rate of 31%, and a recovery of 4 out of 8 actives, or 50%. If we were to use a less stringent approach and include neutral mutations—that is, proceeding with experimental testing of all cases with calculated ΔΔ*G* of +0.5 kcal/mol or less, this would result in experimental measurement of 100 variants, a positive hit rate of 7%, and recovery of 88%, with 7 out of 8 true actives in the experimental dataset classified as neutral or favorable in FEP+ predictions, and leaving only 1 false negative (experimentally active but predicted unfavorable by FEP+), while reducing the number of experimental mutations by 50%, potentially providing significant speedup and cost savings.

### Case studies for application of protein binding FEP+ calculations

We also sought to validate the automated outlier classification scheme above using “real-world” test data. In Figure 6(a), we present the results of running 100 ns protein binding FEP+ calculations with FEP+ Groups treatment of alternate protonation states and automated probable outlier classification and empirical correction, as described above, on a dataset of three case studies from previously active protein biologics design projects. The experimental dataset included SPR measurements for *wt* and single-residue mutation variants for two antibody-antigen systems and one peptide-target system. The antibody-antigen systems included PDB 3SKJ, a complex between the Fab region of the 1C1 antibody and EphA2, with SPR binding affinity measurements for the *wt* complex and 18 Fab 1C1 mutations (presented in Supplemental Table 1); and PDB 5L6Y, a complex between the Fab region of tralokinumab and IL-13 antigen, with measurements for *wt* plus 6 tralokinumab variants (Supplemental Table 2). The cyclic peptide-target complex, presented in Supplemental Table 3, were based on PDB structure 6SBA, a complex between VGL4-mimetic cyclic peptide 4E and mouse TEAD, and included the *wt* complex and 18 peptide variants, including several measurements from a previous study [47]. The experimental binding data, including kinetics measurements for the two antibody systems, are also available in machine-readable form in the GitHub repository accompanying this study (see Data Availability).

**Figure 6:**
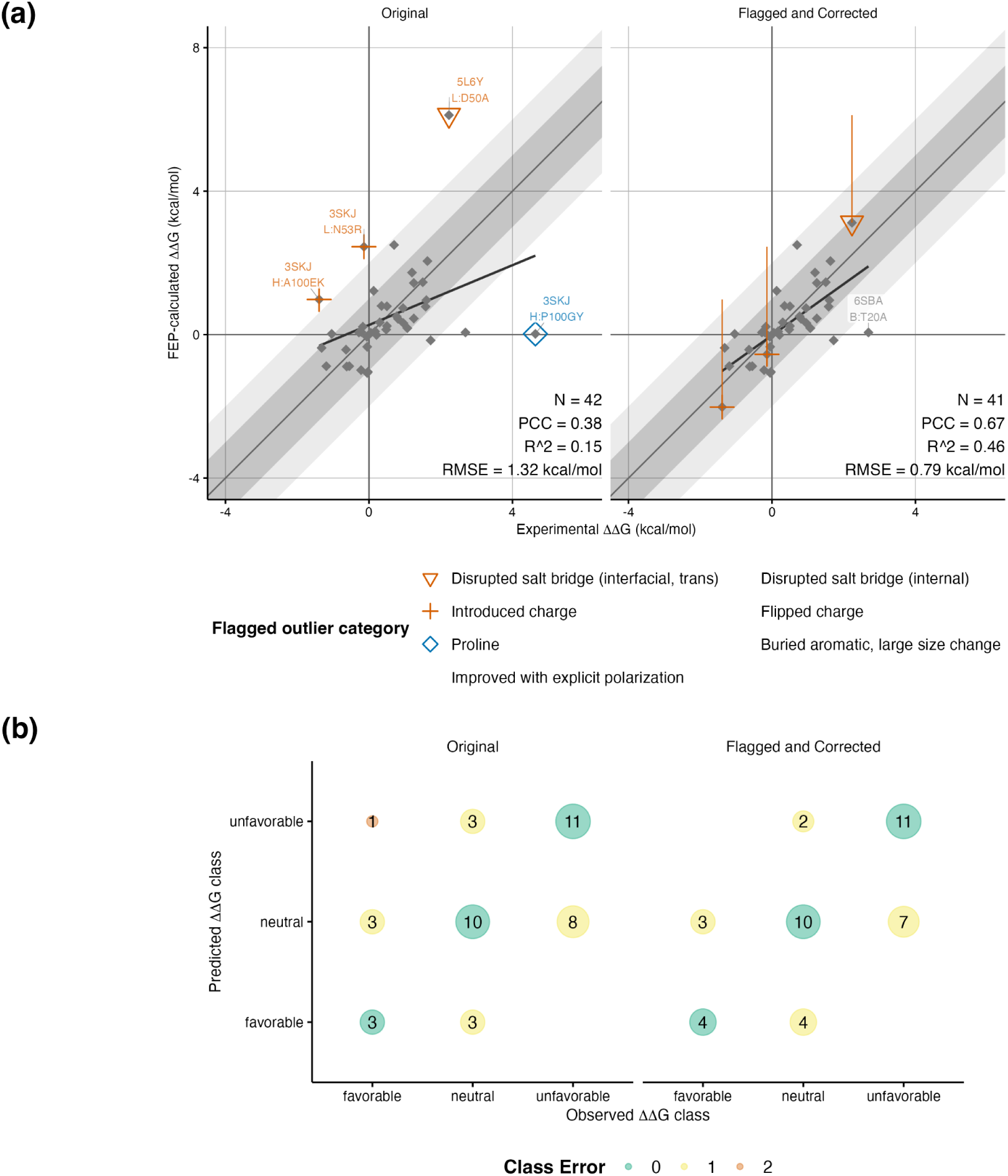
Application of automated outlier flagging and correction to the case studies dataset. (a) Correlation plots and (b) three-class confusion matrices for the 100 ns FEP+ Groups-treated results vs. experiment, before and after automated outlier flagging and empirical correction of charged outlier cases as in Figure 5. The magnitude of the empirical correction applied to the flagged charged outlier cases in (a) was determined using the flagged cases from the benchmark dataset only.

The results for the case studies dataset were similar to those from the benchmark set, with initial RMSE of 1.32, though due to a large proline mutation outlier, a relatively low initial R^2^ of 0.15 for the full FEP+ Groups-treated 100 ns dataset. The effect of the automated outlier flagging and empirical correction procedure was also similar, with 4 out of 5 outlier cases being automatically flagged and 3 of those empirically corrected. After empirical correction of the charged outlier cases and exclusion of the flagged proline mutation case, RMSE was improved to 0.79 kcal/mol, and R^2^ increased substantially to 0.46.

After applying the empirical correction for the charged outlier (using the value fit to the benchmark dataset only) and excluding other automatically flagged cases, only 1 outlier case remained. This remaining outlier, 6SBA B:T20A (expt. 2.88, pred. 0.05), was located at a position on the 6SBA cyclic peptide which, like 1JRH I:E55A in the benchmark dataset, did not interact directly with the target protein in the crystal structure. The large unfavorable experimental binding ΔΔ*G*, in this case, suggests that—as previously hypothesized [47]—a substantial rearrangement occurs for the unbound mutant peptide in the physical system, which was not sampled in simulation, and which would incur a reorganization penalty to adopt the binding-competent conformation present in the complex structure and FEP+ calculations. This is another instance where for mutation of a non-contact residue to have non-negligible effect on binding affinity there must necessarily be a substantive change in conformation. This sort of conformational change can be generally thought to be beyond the domain of applicability of FEP.

As with the benchmark dataset, we achieve good accuracy in three-class classification of the case studies results, as depicted in Figure 6(b). Across classification results for the three systems, overall balanced accuracy was 0.70, and after outlier flagging and empirical correction, no cases resulted in a classification error larger than 1 class. Treated collectively as a hypothetical design project, using the same strict cutoff criteria as described in the previous section, this classification would result in the experimental testing of only 8 variants, with a positive hit rate of 50%, and a recovery of 4 out of 7 actives, or 57%. Using the less stringent approach and including neutral mutations would result in experimental measurement of 28 variants, a positive hit rate of 25%, and recovery of 100%, with 7 out of 7 true actives in the experimental subset corresponding to neutral or favorable FEP+ predictions, and leaving no false negatives, while reducing the number of experimental mutations by 32%.

## Discussion

Here, we have presented results from protein binding FEP+ calculations for a benchmark dataset with a variety of systems. We demonstrated how the overall results were improved using FEP+ Groups treatment to account for the effect of pH and alternate protonation states on binding, and how potentially problematic results were identified, and in some cases empirically corrected, based on chemical, structural, and energetic features from the FEP+ simulations. After the automated flagging and empirical correction, and removal of the flagged probable outlier cases (for which FEP+ results using default protocols were identified as likely to yield ΔΔ*G* results with large errors), for the remaining high-confidence set of 186 cases FEP+ predictions agreed well with experiment, with an RMSE of 1.03 kcal/mol and R^2^ of 0.4. These results are consistent with previous protein FEP+ results where accuracy has been at or near the 1 kcal/mol level[17, 18, 36, 35, 39, 40]. This level of accuracy is approaching the intrinsic uncertainty expected from experimental results, which has been proposed to be in the range of 0.4-0.9 kcal/mol for protein-protein and protein-ligand SPR binding measurements [34, 48, 39].

Some of the largest errors in the full benchmark dataset arose for cases where conformational rearrangement was either observed (as seen with the change in conformation of 1DVF *wt* residue D:H33) or expected (e.g. when mutating to or from proline in a surface loop, or when mutating a solvent-exposed polar residue to hydrophobic Ala in a small cyclic peptide). Cases where substantial rearrangement is required in the physical system may be considered beyond the domain of applicability for protein binding FEP+ calculations run with default protocol. However, we note that if such cases—including the relevant conformations involved—can be identified, potential solutions exist to account for the energetics of well-defined conformational changes. Specifically, protein reorganization FEP has been applied successfully to account for changes in loop conformation in absolute binding FEP calculations with protein-ligand complexes [38]. While the application of protein reorganization FEP to the outlier cases in the benchmark and case studies datasets is beyond the scope of the current study, further investigation of the potential combination of protein binding and protein reorganization FEP may prove valuable.

Perturbations involving proline in particular warrant additional discussion. Proline mutations have been treated before in protein stability FEP+ with reasonable success [35]. However, we have found (both in the current work and in our previous experience with other systems not discussed here) that protein binding FEP+ predictions for proline are significantly more challenging. One reason may be that the solvent and complex legs of binding FEP calculations necessitate two different protein contexts, whereas stability calculations have a linear peptide in the unbound solvent leg with far fewer degrees of freedom and therefore a greater likelihood of achieving adequate sampling of conformational space, similarly to the unbound ligand in a small molecule FEP calculation. Proline perturbations in flexible loops are particularly challenging, as they are prone to adopting different conformations from those preferred by other less torsionally constrained residues. We note that both proline cases in the datasets considered here (one each in the benchmark and case studies sets) were mutations to or from proline in flexible loops at the binding interface.

In addition to sampling errors, we found disruption of a π interaction likely contributed to at least one outlier case in the benchmark set. The current default OPLS4 force field used in FEP+ calculations uses fixed charges only. This leaves it unable to account for π-based interactions, which result from induced polarization upon movement of electronic negative charge relative to the atomic nuclei. The effect here is twofold: first, the energetics of the interactions do not contribute to the energies reported by FEP+ calculations; and second, the lack of explicit stabilization of these interactions in simulation could lead to increased sampling of less relevant conformations, thereby adding energetic noise to the results. Our analysis of the 1JRH I:N53A outlier case, which disrupts a well-defined π-hydrogen bond (Figure 4(e)), demonstrated marked improvement with a developmental force field with explicit polarization. Rigorous validation of a force field with explicit polarization for aromatic and other residue types with delocalized π electrons may help to eliminate some of these kinds of errors, by better accounting for the energetics of π interactions.

The ultimate goal for FEP+ calculations in many use cases is to help guide and ideally streamline and accelerate the experimental component of a molecular design project. While it is possible retrospectively to modify protocols and perform extensive investigations of sampling, starting conformations, and other factors that may affect the accuracy of an FEP-predicted binding ΔΔ*G* result, these sorts of analyses are far more difficult to justify in the absence of data showing some deviation from experiment in the initial prediction. The automatic probable outlier flagging and empirical correction script may serve as a means of prioritizing FEP+ results, such that unflagged or corrected cases may be treated as higher confidence and used more directly as input to a downstream experimental validation pipeline.

It is also prudent to consider that although applying the empirical correction for charged cases does improve accuracy, in most practical biologics optimization use cases, it would be undesirable to leave an unpaired charge buried at the binding interface, and from a design standpoint, there is little difference between an unfavorable predicted ΔΔ*G* of, say, +5 kcal/mol and the corresponding empirically corrected ΔΔ*G* of approximately +2 kcal/mol—neither value would be likely to be pursued experimentally. Finally, we acknowledge that the current probable outlier flagging mechanism, and ongoing development of the method in general, would benefit from a larger dataset with a distribution of mutations less skewed toward alanine, and enough outlier cases of each error class available to allow for a statistically rigorous test/train process, complete with cross-validation, though this is also beyond the scope of the current study.

In summary, the results presented here demonstrate that, for a majority of mutations from a variety of systems, protein binding FEP+ calculations are able to produce accurate results and may provide insight suitable for guiding the experimental component of biologics design projects.

## Methods

### Benchmark dataset curation

Systems were selected from the SKEMPI database to use for FEP+ benchmarking which had a minimum of 20 mutations available for study with ITC or SPR experimental binding measurements; resolution of 2.8 Å or better; and no missing residues or disordered loops in the structure near the experimental mutation sites. Three additional systems from the authors’ previous work were added to the dataset, each with more than 10 mutations with SPR binding measurements and similarly reliable structural coordinates.

For comparison to FEP-calculated binding free energies, experimental binding affinities (𝐾_𝐷_) were converted to Δ*G* values (in kcal/mol) via the standard relation

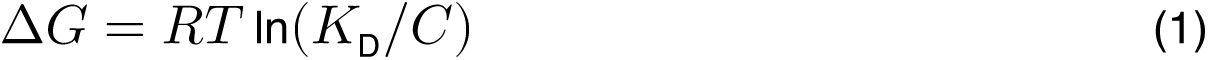

where 𝑅 is the gas constant, 𝑇 is experimental temperature, and 𝐶 is the standard reference concentration of 1 M; and ΔΔ*G* values were determined relative to the *wt* complexes after averaging all available Δ*G* values for each variant.

### Model preparation

Initial structures were obtained from the Protein Data Bank using the accession numbers listed in Table 1. All-atom models were produced using the Protein Preparation Workflow in Maestro (Schrödinger, Inc.), assigning initial protonation states at the experimental pH for each set of measurements using PROPKA [49]. Protein chain termini where the model was truncated compared to the experimental system were capped using acetyl (`ACE`) and N-methylamide (`NMA`) monomers, for the amino- and carboxyl-termini, respectively. Alternate conformations were removed, keeping the highest-occupancy conformation, or the first conformation listed when two conformers were tied for highest occupancy; and any atom clashes were resolved by local minimization. Models were exported to Maestro format (`.mae`) and are available in the GitHub repository associated with this manuscript (see Data Availability).

### Free energy perturbation calculations

FEP calculations were performed using FEP+ (Schrödinger Suites, 2022-2 release), and run with a production stage of 100 ns for each mutation in the assembled dataset, including all possible alternate protonation states for both *wt* and mutant residues using default protocols. Briefly, a graph of mutations was prepared to include all possible direct mutations from the `WT` node (direct perturbations between proline and charged residues are not supported) as well as connecting edges to allow cycle-closure analysis as described previously [44]. For each perturbation edge, complex (bound) and solvent (unbound) simulation legs were run, the former comprising the full prepared complex, and the latter only the component which contained the mutated residue, which in some cases included more than one protein chain. For each leg, the model was placed into an orthorhombic SPC water box with either a 5 Å buffer for neutral and core-hopping (proline) perturbations, or 10 Å buffer for charged perturbations. Sodium or chloride ions were added to neutralize the system, plus additional NaCl ions to 150 mM total concentration for charged perturbations.

Each perturbation leg was run using 12, 16, or 24 lambda windows for neutral, core-hopping, or charged mutations, respectively. Charged perturbations utilized an alchemical ion procedure to maintain system neutrality across all lambda windows, as previously described [18]. Replicas were initialized with the default series of minimization and restrained relaxation steps including a Grand Canonical Monte Carlo (GCMC) solvation protocol in the region around the mutating residue [24], followed by a 100 ns lambda-hopping production stage using the replica exchange with solute tempering (REST) protocol described previously to enhance sampling [23]. All FEP+ calculations used the default OPLS4 force field, except in the analysis of one outlier case, as described in the Results, where a developmental version of a force field with explicit polarization was used. A final analysis stage for each leg included relative Δ*G* calculations using the multistate Bennett acceptance ratio (MBAR) method [50] along with other automated tasks such as trajectory alignment and parching.

### FEP+ Groups treatment of alternate protonation states

To account for the potential shift in pKa of titratable residues upon protein-protein binding, we developed a protocol to be applied to the output FEP+ results for each system. We have adapted the method from [51] for protein systems, using literature micro pKas for titratable residues, along with ΔΔ*G* values between the physical states. Initially, FEP perturbations between `NMA`- and `ACE`-capped single amino acid residues, which represent the “model” physical state, were run in triplicate for all possible amino acid permutations permitted by FEP+, including those not involving titratable residues, and in both directions for each pair of residues (e.g. `ALA->ASP` as well as `ASP->ALA`). We manually compiled these results into CSV format with 4 columns: the protonation state-specific start and end 3-letter residue codes, mean model Δ*G* for the perturbation, and standard deviation among the 3 independent calculations. Typical standard deviations were on the order of 0.02-0.03 kcal/mol, indicating good convergence among the replicates.

To apply the FEP+ Groups treatment to each system, the output FEP+ perturbation map (`.fmp`) file, the pH value of the experimental measurements, and the aforementioned CSV file with precalculated model leg Δ*G* values were provided as input to the `protein_fep_groups.py` script available in the Schrödinger Suites 2024-2 release, and the resulting grouped Δ*G* and pKa values were written to CSV output file. Briefly, this script reads the FEP+ graph; loads the applicable model Δ*G* from the CSV into each perturbation edge; and applies a cycle closure correction to minimize hysteresis around closed thermodynamic cycles. The script aggregates nodes that differ only by protonation or tautomeric state into node groups and reports an FEP+ Groups predicted binding Δ*G* value for each node, which is the same for each node in the node group.

We calculate fractional populations 𝑝 for each node 𝑖 in each physical state for the node group using the following equation:

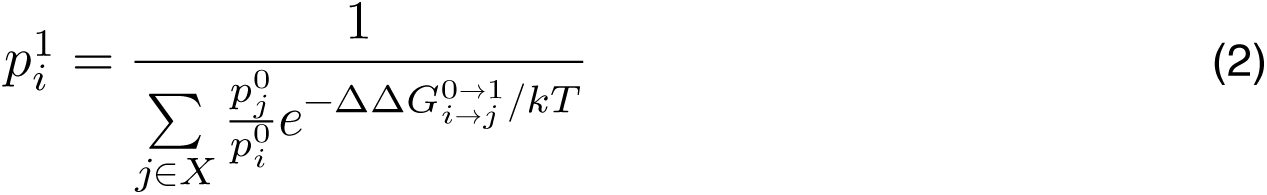

where 𝑋 is the physical ensemble of states represented by the nodes 𝑗 of the node group (including 𝑖), 𝑝^0^ represents a known starting population, and 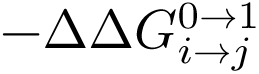 is the FEP-calculated ΔΔ*G* between each pair of nodes 𝑖𝑗 in the node group for the specified physical transition, namely either the model-to-solvent ΔΔ*G*, or solvent-to-complex (binding) ΔΔ*G*. Initial model 𝑝^0^ values for each node in the node group are calculated via the Henderson-Hasselbalch equation:

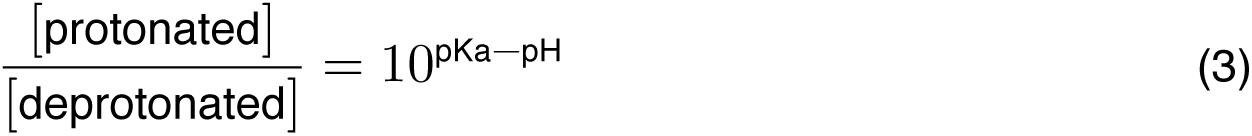

in combination with the differences between precalculated model and solvent leg Δ𝐺 values for each 𝑖𝑗 node pair from the protein binding FEP+ calculation, are used to calculate solvent leg populations 𝑝^1^. The process is repeated using the same formula to calculate complex leg populations (𝑝^2^), using the solvent leg populations 𝑝^1^ and binding 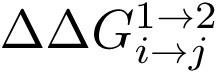 values. After calculating the fractional populations for each node 𝑖, the group predicted binding Δ*G* is calculated by applying state penalties for the complex (2) and solvent (1) states to the original FEP+ predicted Δ*G* value 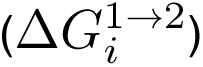 via:

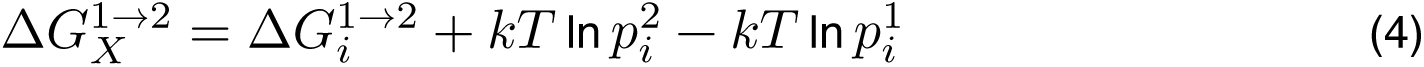

After this FEP groups treatment, all nodes belonging to the same node group—that is, representing the same protein variant—reported identical predicted Δ*G* values, and mutational binding ΔΔ*G* values were calculated for each node group (variant) by subtracting the Protein FEP+ Groups-predicted binding Δ*G* value for the *wt* node group for each system.

### Classification of outliers

To identify limitations in the default protein FEP+ protocol, we individually analyzed cases with calculated binding ΔΔ*G* that deviated from the experimentally determined ΔΔ*G* by more than 2 kcal/mol. We identified several classes of mutations that appeared to be problematic. These were broadly classified as either related to insufficient conformational sampling or due to limitations in the OPLS4 force field, and more specifically into the outlier classes presented in Table 3 and Figure 4.

As an automated solution to identify lower-confidence cases that may require additional attention or potentially be beyond the domain of applicability of FEP+ calculations, we developed two scripts that, based on an output perturbation map file from an FEP+ job, report a list of individual mutations that meet the criteria for one of the outlier categories. The first script extracts a set of chemical and structural features from the models in the perturbation map file. The second script uses the features to identify probable outliers based on empirical observations from the Protein FEP Benchmark dataset, and reports and applies an (optionally user-supplied) empirical correction to charged outlier cases. The source code for both scripts is available in the GitHub repository accompanying this manuscript (see Data Availability).

### Loop prediction

Prediction of loop conformations for two cases on the 1DVF antibody H3 loop were done for the unbound antibody (chains D and E) using the `loop_predictor.py` script available in the Schrödinger Suite, 2024-1 release. For each mutant loop, residues D:94-102, the crystal structure conformation was optimized in-place using the `-repredict_input` command-line flag to allow meaningful comparison of energies. Subsequent conformational searches for each loop were performed with Cα distance constraints of 4, 6, and 8 Å from the crystal structure conformation, and restricting nearby sidechain sampling to include only the residue sidechains sampled in the initial repredict-input calculation. The Prime Energy and loop backbone RMSD were reported for the lowest-energy loop model from among the three conformational searches and compared to those of the lowest-energy loop from the optimized crystal structure conformation.

### Data analysis and figure preparation

Raw FEP data and structural features were extracted and compiled using Python [52] and the Schrödinger Python API, and downstream analyses were done using either Python or R and RStudio [53]. Plots were made using `ggplot2` R package [54]. Structural figures were created using PyMOL [55]. The manuscript was prepared using Quarto and RStudio [56, 57].

## Data availability

Experimental data, prepared all-atom structural models, and all code needed to run the the probable outlier flagging protocol are available in the GitHub repository accompanying this manuscript (https://github.com/schrodinger/protein_fep_benchmark).

## CRediT authorship contribution statement

**Jared M. Sampson:** Conceptualization, Methodology, Software, Formal analysis, Investigation, Data Curation, Writing - Original Draft, Writing - Editing & Review, Visualization. **Daniel A. Cannon:** Methodology, Investigation, Data Curation, Writing - Editing & Review. **Jianxin Duan:** Methodology, Investigation, Data Curation, Writing - Editing & Review. **Alina P. Sergeeva:** Investigation, Data Curation, Writing - Editing & Review. **Jordan C. K. Epstein:** Software, Writing - Editing & Review. **Phinikoula Katsamba:** Investigation, Writing - Editing & Review, Visualization. **Seetha M. Mannepalli:** Investigation, Writing - Editing & Review. **Fabiana A. Bahna:** Investigation, Writing - Editing & Review. **Hélène Adihou:** Conceptualization, Methodology, Validation, Formal analysis, Writing - Editing & Review. **Stéphanie M. Guéret:** Conceptualization, Methodology, Validation, Formal analysis, Writing - Editing & Review. **Ranganath Gopalakrishnan:** Conceptualization, Methodology, Validation, Formal analysis, Writing - Editing & Review. **Stefan Geschwindner:** Methodology, Validation, Formal analysis, Resources, Writing - Editing & Review. **Leonardo De Maria:** Conceptualization, Resources, Data Curation, Writing - Editing & Review, Project administration. **Juan Carlos Mobarec:** Validation, Investigation, Writing - Editing & Review, Supervision, Project administration. **Roger B. Dodd:** Formal analysis, Investigation, Writing - Editing & Review, Supervision. **D. Gareth Rees:** Formal analysis, Investigation, Writing - Editing & Review, Visualization. **Anna Sigurdardottir:** Formal analysis, Investigation, Writing - Editing & Review, Visualization. **Trevor Wilkinson:** Validation, Investigation, Resources, Writing - Editing & Review. **Andrew Buchanan:** Conceptualization, Investigation, Resources, Data Curation, Writing - Editing & Review, Supervision, Project administration, Funding acquisition. **Lawrence Shapiro:** Conceptualization, Resources, Writing - Editing & Review, Supervision, Project administration, Funding acquisition. **Barry Honig:** Conceptualization, Methodology, Resources, Writing - Editing & Review, Supervision, Project administration, Funding acquisition. **Richard A. Friesner:** Conceptualization, Methodology, Investigation, Resources, Writing - Editing & Review, Supervision, Project administration, Funding acquisition. **Lingle Wang:** Conceptualization, Methodology, Investigation, Resources, Writing - Editing & Review, Supervision, Project administration, Funding acquisition.

## Acknowledgements

The authors thank Gregory A. Ross for helpful discussions about group corrections; Addison Schile for assistance in implementing the Protein FEP+ Groups script; Edward Miller for providing guidance with outlier classification code and loop modeling; and Delaram Ghoreishi, Wolfgang Damm, and Edward Harder for assistance running preliminary calculations with explicit polarization.

The work was supported by the Bill and Melinda Gates Foundation, INV-016167 (B.H., L.S., and R.A.F.) and the NIH grant R35-GM139585 (B.H.).

## Supplemental Information

### Supplemental Methods

#### Experimental methods for the DIP/Dpr system

For the 5EO9 and 6WNA systems, *Drosophila melanogaster* defective proboscis extension response (Dpr) and Dpr-interacting protein (DIP) proteins were produced as previously described [42]. Briefly, DNA sequences encoding DIP and Dpr extracellular regions were amplified by PCR and sub-cloned into the mammalian expression vector VRC-8400, modified with a N-terminal BiP signal peptide (BiP: MKLSLVAAMLLLLSAARA) and a C-terminal hexa-histidine tag. Point mutations were introduced into the constructs using the KOD hot start polymerase (Novagen) following the standard QuikChange protocol (Agilent). The constructs were expressed in suspension-adapted HEK293 Freestyle cells (Invitrogen) in serum-free media using polyethylenimine as a transfectant (Polysciences). Growth media was harvested 5 days after transfection, and the secreted proteins were purified by nickel affinity chromatography followed by size exclusion chromatography. Most proteins were stored in a buffer of 10 mM Bis–Tris pH 6.6 and 150 mM NaCl. DIP-α and its mutants were stored in a modified buffer (10 mM Bis–Tris pH 6.0 and 150 mM NaCl) to improve stability. UV absorbance at 280 nm was used to determine protein concentration and verification of purity was determined by gel electrophoresis.

SPR binding assays for DIP/Dpr binding were performed using a Biacore T100 biosensor equipped with a Series S CM4 sensor chip. DIP-α and DIP-α M131F were immobilized over independent flow cells using amine-coupling chemistry in HBS-P pH 7.4 (10 mM HEPES, 150 mM NaCl) buffer at 25°C as described by the manufacturer (Cytiva), resulting in typical immobilization levels of 700-900 resonance units (RU). BSA was immobilized on the reference flow cell.

Binding analysis was performed at 25°C in a running buffer of 10 mM Tris-HCl, pH 7.2, 150 mM NaCl, 1 mM EDTA, 1 mg/ml BSA and 0.01% (v/v) Tween-20. Dpr analytes were prepared in running buffer using a three-fold dilution series and tested at the following concentration ranges: 1) Dpr10 A86M, 81-0.0123 μM, 2) Dpr10 H90A, 27-0.0123 μM, 3) Dpr10 H94A, 9-0.0014 μM, 4) Dpr10 S146M, 27-0.0123 μM, 5) Dpr10, 27-0.004 μM. In each experiment, every concentration range was tested in duplicate. Dpr association phase was monitored for either 30 or 40 s, followed by 120 s dissociation phase, each at 50 μl/min. A buffer wash at 100 μl/min for 60 s was performed at the end of the binding cycle. The analyte was replaced by buffer every three binding cycles to double-reference the binding signals by removing systematic noise and instrument drift. The responses were plotted against the concentration of Dpr and the data was fit to 1:1 interaction model and the 𝐾_𝐷_was calculated as the analyte concentration that would yield half-maximal response. The data was processed using Scrubber 2.0 (BioLogic Software).

#### Experimental methods for the EphA2/mAb 1C1 system

Soluble EphA2 protein was produced as previously described [58]. Fab of 1C1 [59] and its variants were expressed and purified as previously described [60] and monomerized by size exclusion chromatography before use.

SPR binding affinity measurements between Fab and EphA2 were conducted using a Biacore 8K (Cytiva). N-terminal Avi tag biotinylated soluble human EphA2 was titrated on a streptavidin surface prepared on a C1 chip using the standard amine coupling protocols recommended by the manufacturer (Cytiva). Fab was then passed over the immobilized antigen using serial dilutions (400 nM to 3.125 nM including blank controls) with a 180 s association injection, followed by a 600 s dissociation phase, and regeneration with 2 pulses of 3.0 M MgCl_2_ for 20 s, all at a flow rate of 50 μl/min.

#### Experimental methods for the Interleukin-13/Tralokinumab system

Tralokinumab variants were generated, expressed as IgG, and purified as previously described [61]. SPR binding affinities were measured between Tralokinumab and its variants and IL-13 using a Biacore T100 instrument as previously described [61]. During each analysis cycle, after titration of between 42 and 155 RU of IgG onto the protein G′ surface at 5 μl/min, binding to human IL-13 (PeproTech) was assayed at a flow rate of 50 μl/min for a 5 min association step, followed by dissociation for 10-30 min.

#### Experimental methods for peptide synthesis of TEAD peptides

Peptides were synthesized on a Syro I peptide synthesizer (Multisyntech) following standard Fmoc-protocols for solid-phase peptide synthesis. Peptide synthesis was performed on Rink amide LL AM/MBHA resin (Merck, 0.28 mmol/g, 0.05 mmol). To prepare the solutions, the amino acids were dissolved in a solution of oxyma pur 0.5 M in DMF, to obtain a final concentration of 0.5 M, HATU was dissolved in DMF to get a concentration of 0.5 M, DIPEA was dissolved in NMP to get a concentration of 2 M, piperidine was dissolved in DMF with a concentration of 25% and acetic anhydride was dissolved in NMP with a concentration of 10%. For the coupling, 400 μl of amino acids solution (0.2 mmol, 4 eq.) was mixed with 400 μl of HATU solution (0.2 mmol, 4 eq.) and 200 μl of DIPEA solution (0.4 mmol, 8 eq.) and added to the resin for 40 min. The amino acid couplings were followed by a capping of the remaining free amino groups with 800 μl of Ac_2_O solution in presence of 200 μl of DIPEA solution for 2 min. Fmoc-deprotection was performed with 25% piperidine in DMF for 10 min. The subsequent aspartic acid and threonine residues in the sequence were introduced as a pseudoproline building block (Cas no. 920519-32-0). The resulting peptide resin was washed six times with 2 ml of DMF between each step. Each amino acid was coupled twice or thrice.

#### Experimental methods for the mTEAD4/Peptide 4E system

Soluble mouse TEAD4 (mTEAD, residues 210-427) was expressed and purified, and mutational variants of cyclized peptide 4E were synthesized and purified as previously described [47].

SPR experiments were performed on a Biacore S200 unit as previously described [47], with mTEAD4 tethered to the biosensor dextran surface, and serial dilutions of cyclized peptide analyte in 0.3% (v/v) DMSO, typically starting from a peptide concentration of 30 μM. Association time was 45 s, with dissociation monitored for 6 min.

#### Surface plasmon resonance of TEAD peptides

The SPR experiments were either performed on a Biacore S200 optical biosensor unit or a Biacore 8K optical biosensor unit using Series S CM5 (Research grade) sensor chips (Cytiva). Prior to use, the sensor chips were equilibrated at room temperature for 15 min to prevent water condensation on the detector side of the sensor chip surface. A running buffer was prepared composed of 10 mM HEPES, 150 mM NaCl, and 0.05% (w/v) Tween-20, pH 7.4, and the system was equilibrated at 20°C using a flow rate of 30 μl/min after docking of the sensor chip. Ligand binding experiments were performed applying the concept of multi-cycle kinetics. A contact time of 45 s was selected, followed by a 6 min dissociation phase to allow for complete dissociation of the analyte prior to the next cycle. The peptides were dissolved in DMSO to a stock concentration of 10 mM. A digital dispenser (HP D300, Tecan) was used to dispense varying concentration of the ligands into running buffer provided in a standard 384-well plate and normalized with DMSO to 0.3% (v/v). Typically, seven concentrations of the analytes were examined, applying a threefold dilution pattern with 30 μM as the top concentration. For the analysis, five running buffer blanks were injected to equilibrate the instrument. The data collection rate was set to 10 Hz, and all experiments were repeated at least three times to allow for error estimations. The data were analysed using Genedata Screener for SPR using the implemented steady-state data fitting routines and by applying a 1:1 binding model for the estimation of peptide affinities.

#### Surface tethering of GST-hTEAD1 and mTEAD4 for SPR

For the covalent tethering of mTEAD4 onto the CM5 biosensor chip, running buffer at a flow-rate of 10 μl/min was used. The carboxyl-dextran surface was activated for 7 min with 0.05 M NHS and 0.2 M EDC, followed by an injection of the mTEAD protein in 10 mM MES pH 6.4 at a concentration of 30-50 μg/ml. Contact times of 2-3 min were sufficient to achieve the desired densities of 2000-3000 RU. This was followed by a deactivation of the residual esters by injecting a solution of 0.5 M ethanolamine pH 8.0 for 7 min before engaging in ligand binding experiments. Reference surfaces were prepared accordingly, omitting the injection of protein over the activated reference surface.

#### Supplemental Figures

**Supplemental Figure 1:**
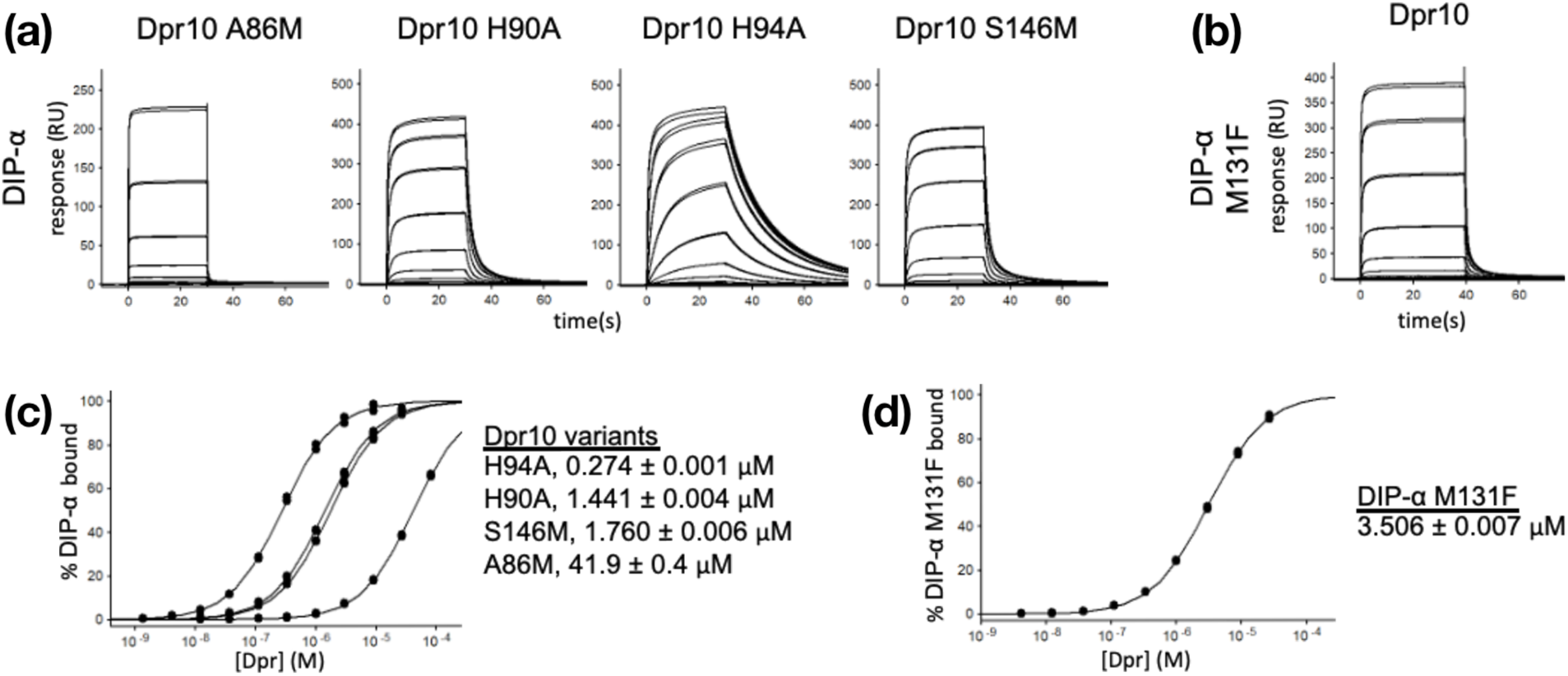
SPR measurements for the Dpr10/DIP-α (6NRQ) system. (a-b) Binding traces for (a) Dpr10 mutants A86M, H90A, H94A and S146M binding to DIP-α; and (b) Dpr10 to the DIP-α M131F mutant. (c-d) Binding isotherms from the experiments in (a) and (b) were used to calculate 𝐾_D_ values for the (c) Dpr10 and (d) DIP-α mutants, respectively, which are shown next to each binding curve with experimental fitting uncertainties.

**Supplemental Figure 2:**
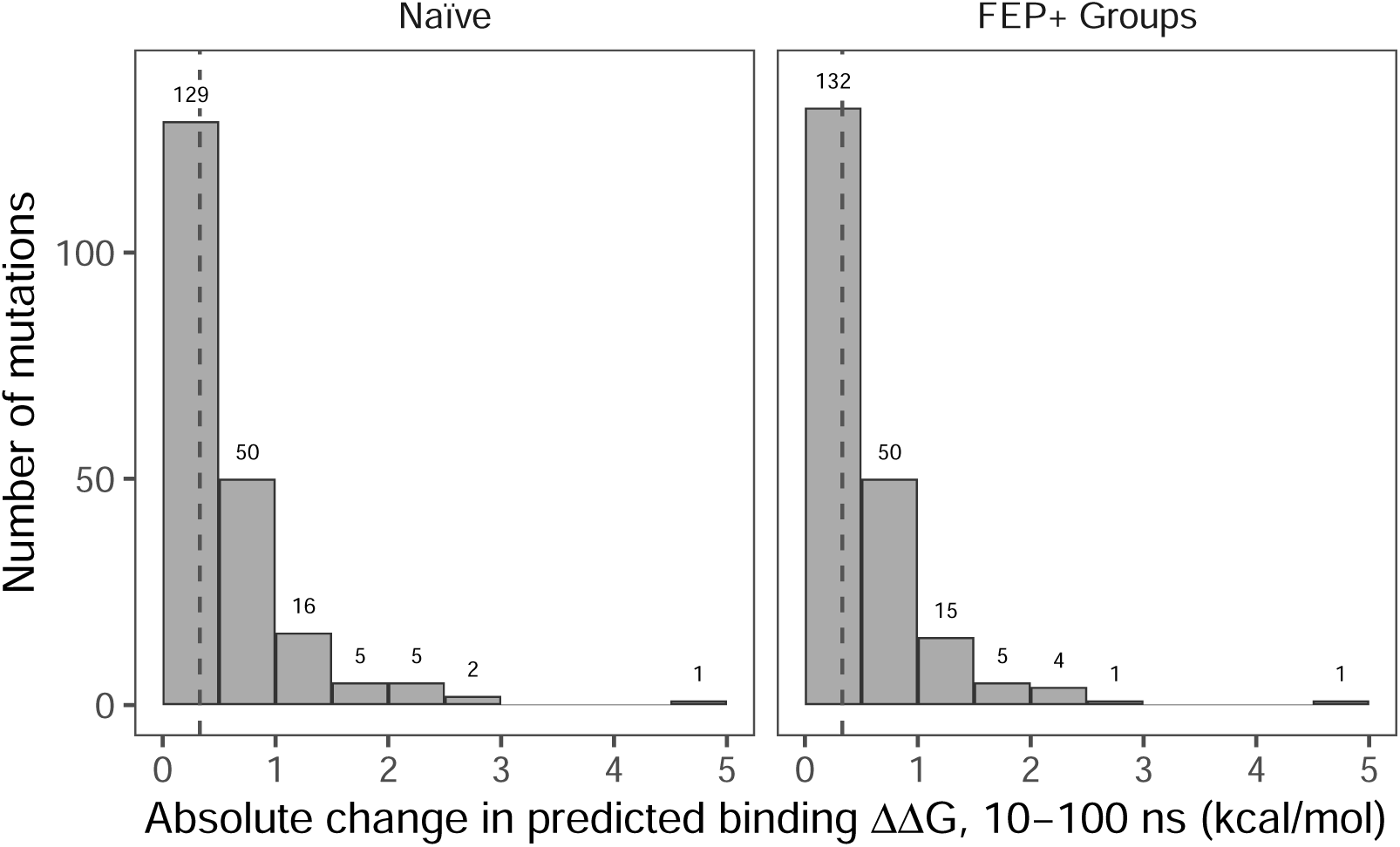
Summary of absolute shifts in predicted binding ΔΔ*G* between 10 and 100 ns timepoints for Naïve and FEP+ Groups-treated results for the benchmark dataset. The number of cases is shown for each bin, and median ΔΔ*G* values of 0.33 and 0.33 kcal/mol, respectively, are indicated by vertical dashed lines. The vast majority of cases (86 and 88% for naïve and FEP+ Groups-treated results, respectively) produced ΔΔ*G* values at 10 ns that were within 1 kcal/mol of the corresponding value at 100 ns.

**Supplemental Figure 3:**
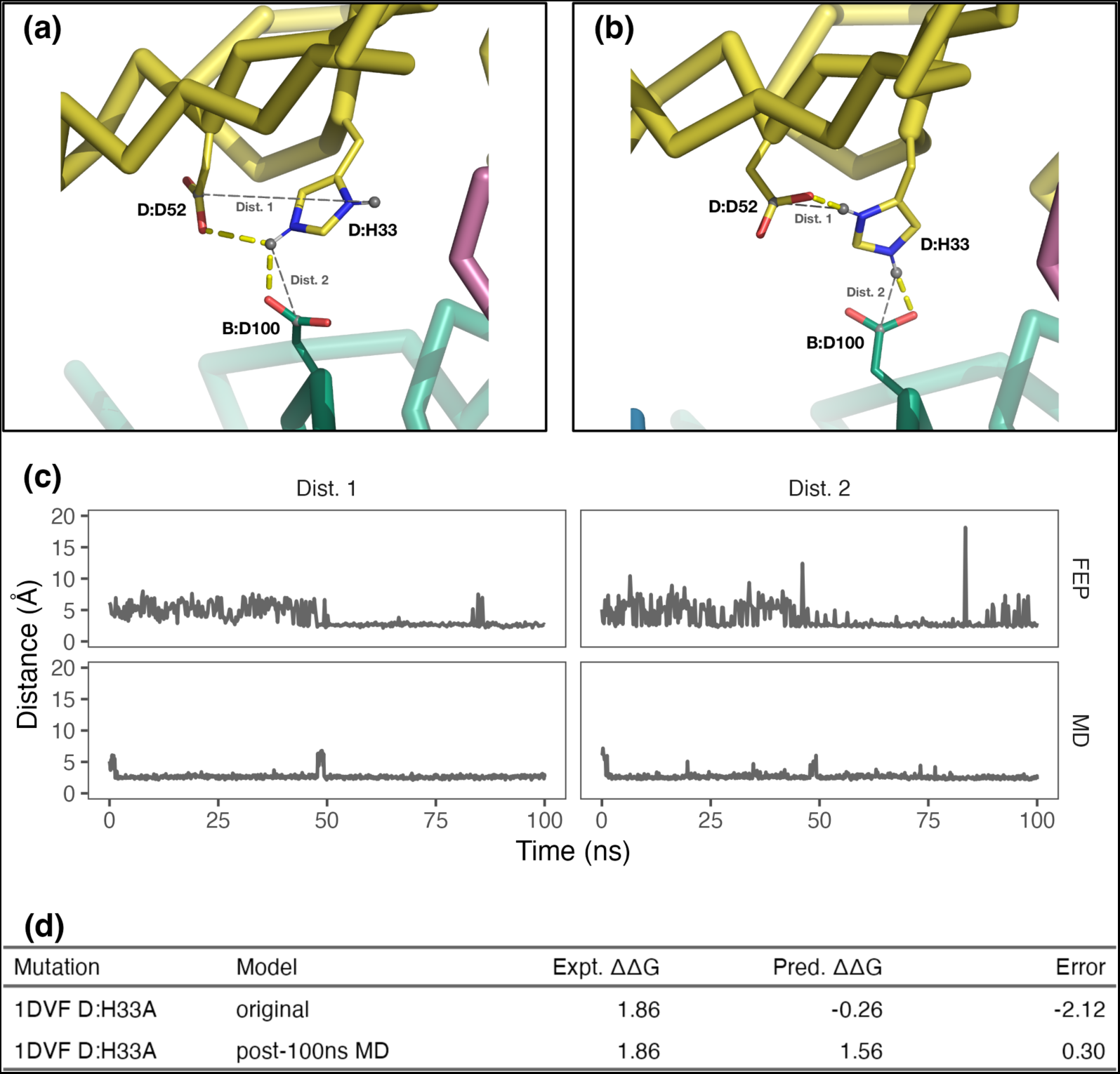
Analysis of the 1DVF D:H33A outlier case. (a) The starting conformation of D:H33 in the prepared 1DVF crystal structure was suboptimal, with a single H33 side chain N-H group interacting with both carboxylates. (b) During 100 ns standard MD, these three side chains quickly relaxed to form two salt bridges, shown here in the configuration present in the final frame of the MD trajectory. Distance measurements 1 and 2 are indicated between `D:HIP33.ND1`—`D:ASP52.CG` (*cis*) and `D:HIP33.NE2`—`B:ASP100.CG` (*trans*), respectively. (c) Distances 1 and 2 as defined in (a) and (b) are shown for the 100 ns MD trajectory and the HIP endpoint trajectory from the `D:HIP33->HID` perturbation edge. The two-salt-bridge conformation in (b) was only sampled after approximately 50 ns in the FEP+ simulation (upper panels), but was maintained nearly exclusively throughout the MD trajectory (lower panels). (d) FEP+ simulation of the D:H33A mutation (including perturbation to alternate His protonation states) using the original starting model produced an absolute error over 2 kcal/mol; but using an MD-relaxed model—namely, the final frame from the 100 ns MD simulation shown in (b)—yielded an accurate prediction with an absolute error of only 0.3 kcal/mol after Protein FEP+ Groups treatment.

**Supplemental Figure 4:**
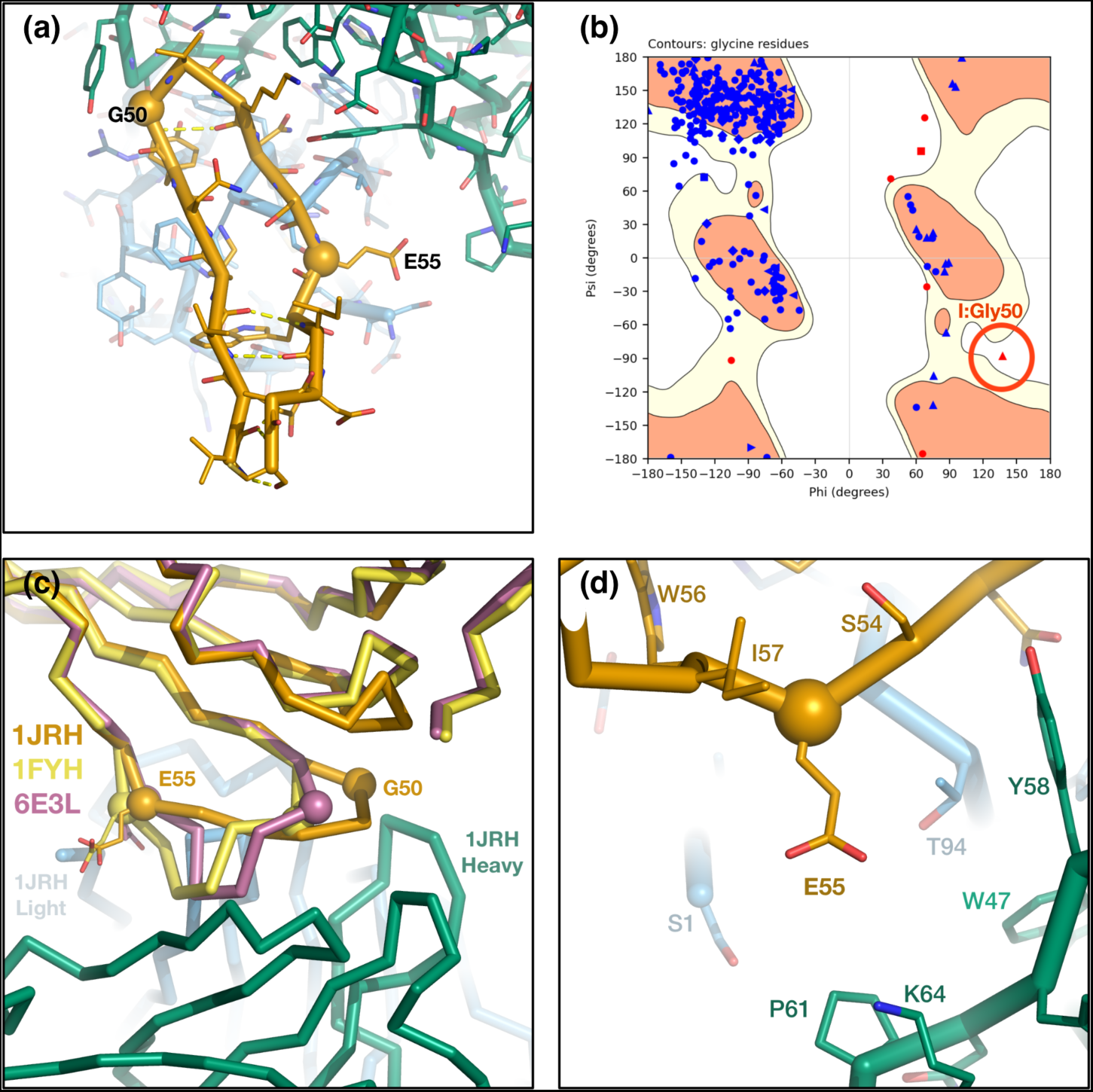
Analysis of the 1JRH I:G50A and I:E55A/P outliers. (a) IFNγR1 residues I:G50 and I:E55 are part of a flexible loop (I:50-56) that, in the 1JRH complex with the A6 Fab fragment, adopts a tight antiparallel structure resembling that of a β-hairpin, but lacks the typical β-strand hydrogen bonding pattern. Here the loop is shown (omitting the rest of the IFNγR1 protein for clarity) in orange with Cα atoms of G50 and E55 as spheres. The Fab A6 heavy and light chains are shown in green and light blue, respectively. (b) Residue I:Gly50 is an outlier in the crystal structure Ramachandran plot, with Phi/Psi angles of 137°/-88°. The presence of a Ramachandran outlier in a flexible loop suggests either an error in model building or crystal packing-induced strain, and in either case represents a high-energy starting conformation, from which relaxation to the true lowest-energy conformation in relatively short FEP timescales is uncertain. (c) Other crystal structures of IFNγR1, including PDB 1FYH chain B (yellow) and 6E3L chain D (violet), are aligned to 1JRH chain I backbone atoms and adopt conformations that are similar to each other, but distinct from the conformation of 1JRH chain I (orange), suggesting a lower energy conformation for the I:50-56 loop in the absence of the antibody and/or 1JRH crystal context. Gly50 and Glu55 Cα atoms are shown as a spheres for each IFNγR1 chain. Sampling between distinct conformations like these is challenging within the relatively short timescales used for FEP+. (d) Wild-type IFNγR1 residue I:E55 does not contact the antibody or form any specific interactions in the 1JRH crystal structure, suggesting any effect of its mutation on binding affinity must involve conformational change of the I:50-56 loop, overall domain orientation, or both.

**Supplemental Figure 5:**
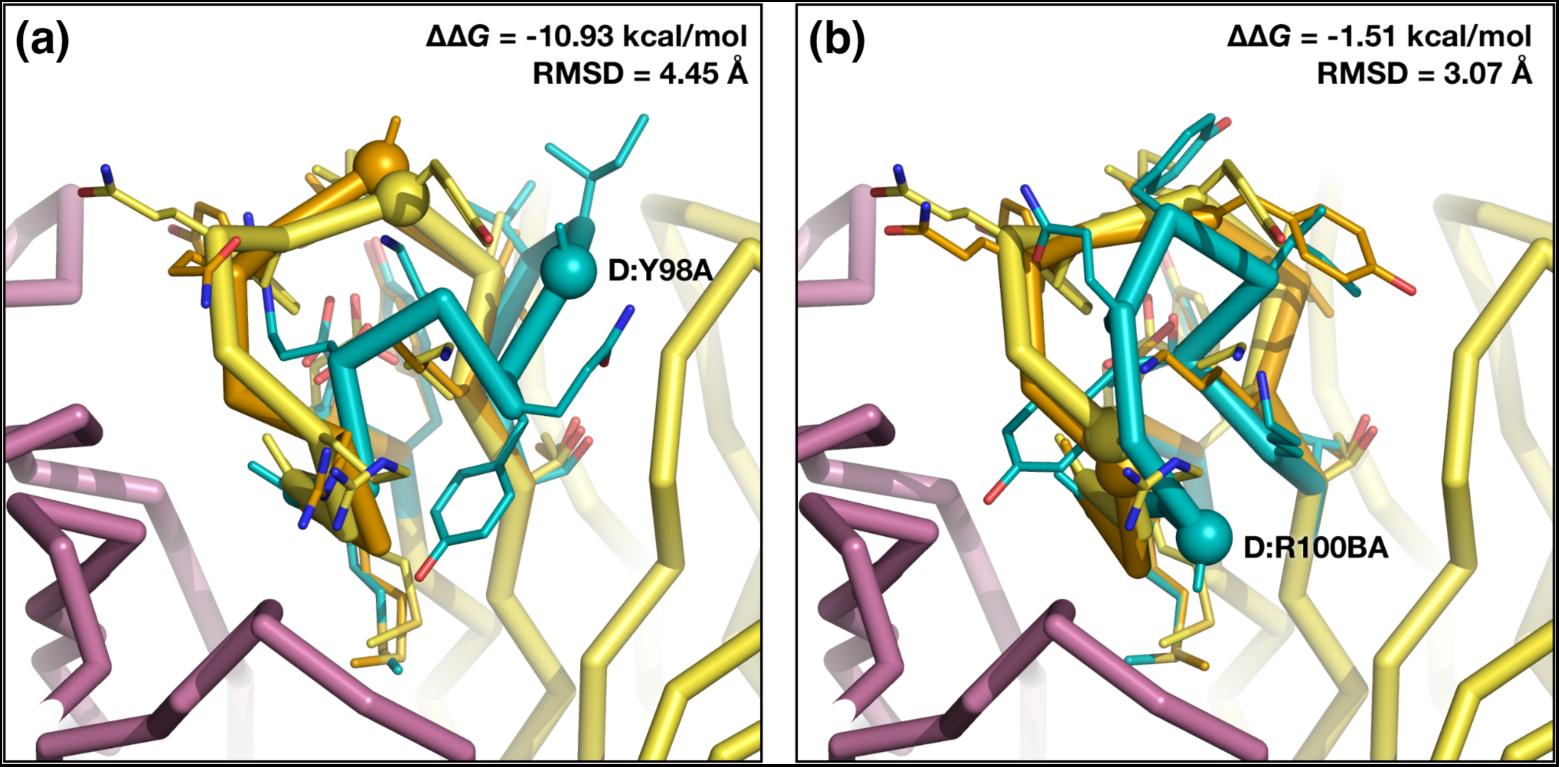
Analysis of the 1DVF D:Y98A/D:R100BA outlier cases. CDR H3 loop conformations for (a) D:Y98A and (b) D:R100BA variants. The crystal structure light and heavy chain variable regions are shown in violet and yellow, respectively. An optimized crystal structure-like conformation is shown in orange, and the lowest energy predicted loop in teal, with Cα atoms of the mutated residues shown as spheres. Backbone RMSD of the lowest energy loop compared to the crystal structure loop, and ΔΔ*G* of the lowest energy loop relative to the repredicted crystal structure conformation are indicated.

#### Supplemental Tables

**Supplemental Table 1:** SPR measurements for the 3SKJ system. Binding kinetics association (𝑘_a_) and dissociation (𝑘_d_) constants and binding affinities (𝐾_D_) are indicated for heavy (H) and light (L) chain Fab 1C1 variants binding to immobilized EphA2 antigen. Mutations are listed using Kabat numbering.

| IgG Variant | $k_a$ (M<sup>-1</sup>s<sup>-1</sup>) | $k_d$ (s<sup>-1</sup>) | $K_D$ (nM) |
| --- | --- | --- | --- |
| WT | 2.67e+05 | 1.60e-02 | 59.6 |
| H:G52R | 3.61e+05 | 1.99e-02 | 55.1 |
| H:A100CI | 1.46e+04 | 1.35e-02 | 929 |
| H:V100DM | 2.82e+05 | 1.05e-02 | 37.3 |
| H:A100EF | 2.22e+05 | 2.41e-01 | 1090 |
| H:A100EI | 2.23e+05 | 6.56e-02 | 294 |
| H:A100EK | 3.17e+05 | 1.80e-03 | 5.67 |
| H:A100EM | 3.41e+05 | 8.01e-03 | 23.5 |
| H:P100GY | 1.61e+03 | 2.42e-01 | 150000 |
| L:S30W | 6.39e+05 | 3.48e-02 | 54.5 |
| L:T31F | 3.05e+05 | 1.26e-02 | 41.3 |
| L:T31H | 2.75e+05 | 1.56e-02 | 56.7 |
| L:T31K | 4.78e+05 | 3.85e-03 | 8.06 |
| L:T31Q | 3.53e+05 | 7.47e-03 | 21.1 |
| L:T31R | 5.20e+05 | 1.05e-02 | 20.2 |
| L:N53R | 4.12e+05 | 1.94e-02 | 47.1 |
| L:N53Y | 2.20e+05 | 4.29e-02 | 195 |
| L:T56F | 2.92e+05 | 1.23e-02 | 42.0 |
| L:T56W | 2.70e+05 | 1.21e-02 | 44.7 |

**Supplemental Table 2:** SPR measurements for the 5L6Y system. Binding kinetics association (𝑘_a_) and dissociation (𝑘_d_) constants and binding affinities (𝐾_D_) are indicated for IL-13 antigen binding to immobilized heavy (H) and light (L) chain Tralokinumab IgG variants. Mutations are listed using Kabat numbering.

| Fab Variant | $k_a(\text{M}^{-1}\text{s}^{-1})$ | $k_d(\text{s}^{-1})$ | $K_D(\text{pM})$ |
| --- | --- | --- | --- |
| WT | 5.17e+06 | 3.25e-04 | 63 |
| H:S99A | 6.07e+06 | 3.46e-04 | 57 |
| L:I27A | 4.95e+06 | 2.57e-03 | 519 |
| L:I27N | 4.90e+06 | 4.60e-03 | 940 |
| L:D50A | 6.00e+05 | 1.63e-03 | 2700 |
| L:G68E | 5.20e+06 | 2.70e-03 | 520 |
| L:G68W | 4.20e+06 | 3.80e-04 | 90 |

**Supplemental Table 3:** SPR measurements for the 6SBA system. Binding affinities (𝐾_D_) are indicated for cyclic peptide 4E variants binding to immobilized mTEAD. Complete peptide sequences (residues 8-27) are shown; macrocyclization was achieved via an amide bond between the Glu10 and Lys25 sidechains in each case.

| Peptide Variant | Sequence | $K_D$ (μM) |
| --- | --- | --- |
| WT | SVEDHFAKALGDTWLQIKAA | $1.2 \pm 0.1$ |
| B:S8A | AVEDHFAKALGDTWLQIKAA | $2.7 \pm 0.25$ |
| B:V9A | SAEDHFAKALGDTWLQIKAA | $1.5 \pm 0.37$ |
| B:D11A | SVEAHFAKALGDTWLQIKAA | $1.6 \pm 0.69$ |
| B:K15A | SVEDHFAAALGDTWLQIKAA | $2.8 \pm 0.06$ |
| B:T20A | SVEDHFAKALGDAWLQIKAA | $113.0 \pm 3.5$ |
| B:W21A | SVEDHFAKALGDTALQIKAA | $15.0 \pm 0.85$ |
| B:L22A | SVEDHFAKALGDTWAQIKAA | $2.0 \pm 0.14$ |
| B:Q23A | SVEDHFAKALGDTWLAIKAA | $1.3 \pm 0.11$ |
| B:I24A | SVEDHFAKALGDTWLQAKAA | $4.4 \pm 1.35$ |
| B:V9F | SFEDHFAKALGDTWLQIKAA | $17.4 \pm 1.1$ |
| B:V9W | SWEDHFAKALGDTWLQIKAA | $7.32 \pm 0.06$ |
| B:V9Y | SYEDHFAKALGDTWLQIKAA | $9.16 \pm 0.25$ |
| B:K15F | SVEDHFAFALGDTWLQIKAA | $2.77 \pm 0.04$ |
| B:K15R | SVEDHFARALGDTWLQIKAA | $0.208 \pm 0.02$ |
| B:K15W | SVEDHFAWALGDTWLQIKAA | $6.03 \pm 0.12$ |
| B:K15Y | SVEDHFAYALGDTWLQIKAA | $4.72 \pm 0.11$ |
| B:A16V | SVEDHFAKVLGDTWLQIKAA | $2.17 \pm 0.007$ |
| B:Q23R | SVEDHFAKALGDTWLRIKAA | $0.13 \pm 0.01$ |

